# Mitochondrial structure and function adaptation in residual triple negative breast cancer cells surviving chemotherapy treatment

**DOI:** 10.1101/2022.02.25.481996

**Authors:** Mokryun L. Baek, Junegoo Lee, Katherine E. Pendleton, Mariah J. Berner, Emily B. Goff, Lin Tan, Sara A. Martinez, Tao Wang, Matthew D. Meyer, Bora Lim, James P. Barrish, Weston Porter, Philip L. Lorenzi, Gloria V. Echeverria

## Abstract

Neoadjuvant chemotherapy (NACT) used for triple negative breast cancer (TNBC) eradicates tumors in approximately 45% of patients. Unfortunately, TNBC patients with substantial residual cancer burden have poor metastasis free and overall survival rates. We previously demonstrated mitochondrial oxidative phosphorylation (OXPHOS) was elevated and was a unique therapeutic dependency of residual TNBC cells surviving NACT. We sought to investigate the mechanism underlying this enhanced reliance on mitochondrial metabolism. Mitochondria are morphologically plastic organelles that cycle between fission and fusion to maintain mitochondrial integrity and metabolic homeostasis. The functional impact of mitochondrial structure on metabolic output is highly context dependent and not understood in TNBC. Several chemotherapy agents are conventionally used for neoadjuvant treatment of TNBC patients. Upon comparing mitochondrial effects of commonly used chemotherapies, we found that DNA-damaging agents increased mitochondrial elongation, mitochondrial content, flux of glucose through the TCA cycle, and OXPHOS, whereas taxanes instead decreased mitochondrial elongation and OXPHOS. Additionally, short protein isoform levels of the mitochondrial inner membrane fusion protein optic atrophy 1 (OPA1) were associated with those observations. Further, we observed heightened OXPHOS, OPA1 protein levels, and mitochondrial elongation in a patient-derived xenograft (PDX) model of residual TNBC. Pharmacologic or genetic disruption of mitochondrial fusion and fission resulted in decreased or increased OXPHOS, respectively, revealing that longer mitochondria favor oxphos in TNBC cells. Using TNBC cell lines and an *in vivo* PDX model of residual TNBC, we found that sequential treatment with DNA-damaging chemotherapy, thus inducing mitochondrial fusion and OXPHOS, followed by MYLS22, a specific inhibitor of OPA1, was able to suppress mitochondrial fusion and OXPHOS and significantly inhibited residual tumor regrowth. Taken together, our findings suggest that TNBC mitochondria can optimize OXPHOS through modulation of mitochondrial structure. This may provide an opportunity to overcome mitochondrial adaptations of chemoresistant TNBC.

## INTRODUCTION

Triple negative breast cancer (TNBC) is characterized by lack of estrogen receptor and progesterone receptor expression and lack of amplification of the human epidermal growth factor receptor. Thus, there are limited targeted therapy options for this subtype which comprises 15-20% of all breast cancers^1^. Standard chemotherapies remain a mainstay treatment for this disease. Unfortunately, approximately 45% of TNBC patients with localized disease treated with neoadjuvant (pre-surgical) chemotherapy (NACT) fail to achieve complete pathologic response (pCR) and harbor substantial residual cancer burden (RCB) at the time of surgery, leading to a high risk of distant metastasis and mortality^2-5^. Recently, combination chemotherapy has been augmented by the addition of PDL1 inhibition with a marginal, but statistically significant, increase in efficacy^6^. Thus, there is an urgent need to delineate mechanisms of residual TNBC cell survival so they can be rationally therapeutically targeted.

By analyzing serial pre- and post-NACT tumor samples from orthotopic patient-derived xenograft (PDX) models and biopsies from women with TNBC, we previously found residual tumors following treatment with the standard front-line NACT regiment, Adriamycin (doxorubicin) plus cyclophosphamide (AC), had heightened mitochondrial oxidative phosphorylation (OXPHOS) relative to treatment-naïve tumors^7^. We demonstrated residual tumors had heightened susceptibility to OXPHOS inhibition using an inhibitor (IACS-010759)^8^ of electron transport chain Complex I. Furthermore, TNBC stem-like cells have been shown to have increased OXPHOS activity and chemoresistance^9^. Induction of OXPHOS following chemotherapy or targeted therapy treatment has also been observed in experimental models of leukemias^10-12^, colon^13,14^, prostate^15^ and non-small cell lung^16,17^ cancers. Thus, targeting mitochondrial metabolism may be a promising avenue to overcome chemotherapy-induced adaptations in TNBC.

Mechanisms underlying the enhanced reliance on OXPHOS in residual TNBC cells that survive chemotherapy are yet unknown. The incredible dynamics of mitochondrial structure were first appreciated over 100 years ago in chick embryo cultures^18^. After biogenesis, mitochondria have a ‘life cycle’ that includes structural changes mediated by fission and fusion, ending with mitophagic elimination of fragmented organelles that have lost membrane potential^19^. This interchange between fission and fusion, known as ‘mitochondrial dynamics’, enables metabolic adaptation and survival. It is ultimately the balance between fission and fusion that dictates mitochondrial structure^20^. Mitochondrial fission results in small, fragmented mitochondria, whereas fusion produces elongated, inter-connected mitochondrial networks^21,22^. Mitochondrial fusion is orchestrated by the partially redundant dynamin-like GTPases Mitofusin 1 and 2 (MFN1, MFN2) which fuse the outer membranes of neighboring mitochondria^23,24^. This is coordinated with fusion of the inner mitochondrial membrane by the dynamin-like GTPase optic atrophy 1 (OPA1). OPA1 orchestrates not only mitochondrial fusion but also cristae organization, ETC super-complex assembly, and mitochondrial genome maintenance^25,26^. OPA1 is encoded in the nucleus and alternatively spliced to produce 8 mRNA isoforms. It is then translated in the cytosol, imported into mitochondria, and cleaved by multi-functional proteases YME1L1 and OMA1^27^. Up to five OPA1 C-terminal protein isoforms are typically visualizable by western blotting. Skipping of alternative exon 5b followed by proteolytic cleavage by OMA1 produces the short ‘E’ isoform C-terminal fragment, required for proper inner mitochondrial membrane structure and ETC complex formation^25,28^. In some stress conditions, OMA1 activity is enhanced, resulting in loss of long OPA1 isoforms and accumulation of short OPA1 isoforms^29^.

Conversely, the dynamin 1-like GTPase (DRP1) oligomerizes at endoplasmic reticulum (ER)-mitochondria contact sites to sever mitochondrial membranes, leading to fission^30^. It was recently appreciated that mitochondrial fission can cause either mitophagic elimination of damaged mitochondria or mitochondrial proliferation^31^. Mitochondrial fusion is generally considered a mechanism through which mitochondria exchange and refresh internal components, which in some cases optimizes mitochondrial functioning^32^. However, the functional impacts of mitochondrial fission and fusion are highly varied among tumor types. Mitochondrial fission, for example, drives opposite metabolic effects in pancreatic cancer and neuroblastoma cells, driving increased or decreased OXPHOS, respectively^33,34^. In various TNBC cell line models, mitochondrial fission has been shown to either drive or inhibit apoptosis^35,36^, and fusion has been shown to promote or inhibit metastatic phenotypes^36,37^. Furthermore, expression of mitochondrial fusion genes has been found to correlate with favorable or poor prognosis in various TNBC cohorts^35,36,38^. However, the functional impact of mitochondrial structure on the metabolic output of TNBC mitochondria is yet unknown.

In the present study, we establish a functional role for mitochondrial structure in the metabolic output of mitochondria in TNBC cells. Further, we find that standard of care DNA-damaging chemotherapies induce mitochondrial fusion in residual cells not killed by treatment, while taxanes can induce mitochondrial fragmentation. This functionally contributed to the higher and lower OXPHOS, respectively, induced by these chemotherapies. We find further evidence for this functional relationship in a patient-derived xenograft (PDX) model of residual TNBC. Lastly, we provide *in vitro* and *in vivo* evidence that therapeutically inhibiting mitochondrial fusion is more efficacious in post-chemotherapy residual tumor cells rather than it is in pre-chemotherapy-treated cells. These findings provide rationale for the development of therapeutic targeting strategies that may overcome mitochondrial adaptations in chemoresistant TNBC.

## RESULTS

### Standard chemotherapies alter mitochondrial metabolism in TNBC cells

To delineate the metabolic effects of chemotherapies on TNBC cells, we treated MDA-MB-231 and SUM159pt human TNBC cells with an IC_50_ dose (Fig. S1A) of conventional DNA-damaging agents (doxorubicin, carboplatin) and microtubule-stabilizing taxanes (paclitaxel, docetaxel) that are commonly used in NACT regimens for TNBC. Following 48 hours of treatment, we analyzed metabolism in the residual surviving cell population. We measured oxygen consumption rate (OCR) with the MitoStress kit on a Seahorse bioanalyzer. In contrast, treatment with DNA-damaging chemotherapy agents caused significant increases in basal and maximal OCR (Fig. 1A-B and S1B-C). On the other hand, treatment with taxanes significantly reduced basal and maximal OCR compared to vehicle-treated cells (Fig. 1A-B and S1B-C). Interestingly, basal ECAR was also elevated by DNA-damaging chemotherapy agents (Fig. 1B and S1C).

**Figure 1.**
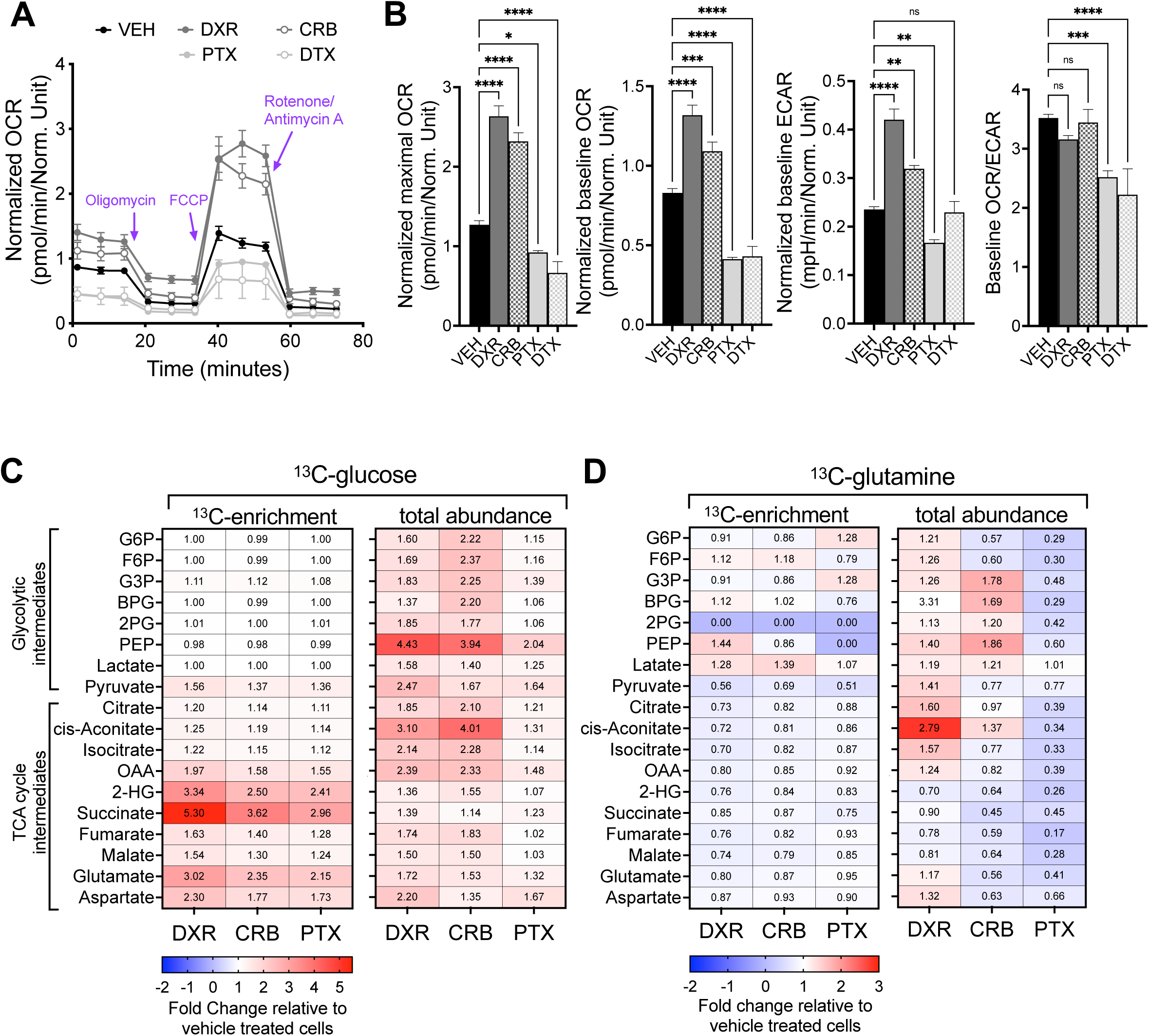
Conventional chemotherapies differentially alter mitochondrial metabolism in TNBC cells. (A) Seahorse Mito Stress assay of MDA-MB-231 cells treated with DNA-damaging chemotherapy agents (DXR, doxorubicin; CRB, carboplatin) and microtubule stabilizing chemotherapy agents (PTX, paclitaxel; DTX, docetaxel) compared to vehicle, with (B) quantified maximal OCR (oxygen consumption rate), basal OCR, basal ECAR (extracellular acidification rate), and OCR/ECAR ratio. Data were normalized to total number of cells stained with DAPI immediately after Seahorse assay *\*\*\*\*P* < 0.0001, ***<0.001, **<0.01, *<0.05 by one-way ANOVA. Data are representative of at least 3 independent experiments. (C-D) ^13^C-glucose (C) derived or ^13^C-glutamine (D) derived-enrichment or total abundance of glycolytic and TCA cycle intermediates in MDA-MB-231 cells treated vehicle, DXR, CRB and PTX. Data were normalized to vehicle treated cells. n=3, G6P (Beta-D-Glucose 6-phosphate), F6P (Fructose 6-phosphate), G3P (Glycerol 3-phosphate), BPG (2,3-Diphosphoglyceric acid), 2PG (2-Phospho-D-glyceric acid), PEP (Phosphoenolpyruvic acid), OAA (Oxoglutaric acid), 2-HG (2-Hydroxyglutarate)

To directly compare relative cytosolic (glycolysis) and mitochondrial (tricarboxylic acid, TCA, cycle) metabolic rates in these *in vitro* models, we conducted stable isotope tracing of energy substrates glucose and glutamine by growing MDA-MB-231 and SUM159pt cells in the presence of ^13^C_6_ -glucose or ^13^C_5_ -glutamine following treatment with an IC_50_ dose of chemotherapy. We used ion chromatography–mass spectrometry (IC-MS) to measure levels of metabolite isotopologs. Although we observed no significant changes in glucose-derived ^13^C labeling of glycolytic intermediates, there was a substantial increase in labeling of multiple TCA cycle intermediates following treatment (Fig. 1C, S1D) in both cell lines. In contrast, we observed no increase in glutamine-derived ^13^C labeling of TCA cycle intermediates following any chemotherapy treatment (Fig. 1D, S1E). These results suggest that chemotherapy treatment induces glucose-fueled TCA cycling and OXPHOS as an adaptive mechanism. Consistent with the increased ECAR observed in chemotherapy-treated cells (Fig. 1B and S1C), we observed increased total abundance of several glycolytic intermediates in chemotherapy-treated cells. Since glycolytic intermediates were increased after chemotherapy treatment but were not derived from labeled glucose, these findings suggest upregulation of an alternative pathway such as lactate-fueled gluconeogenesis after chemotherapy treatment. In summary, we found that DNA-damaging agents increased, whereas taxanes decreased, OXPHOS rate. Of note, DNA-damaging agents and taxanes both induced glucose-driven mitochondrial TCA cycling.

### Changes in mitochondrial metabolism are associated with mitochondrial structure in residual TNBC cells following chemotherapy treatment

To investigate the impact of chemotherapy on mitochondrial structure, we used fluorescent microscopy to inspect mitochondrial shape in MitoTracker-stained residual MDA-MB-231 and SUM159pt cells that survived chemotherapy treatment. While DNA-damaging chemotherapies increased mitochondrial elongation, taxanes either had little effect or diminished elongation (Fig 2A-B, S2A) relative to vehicle-treated cells as evidenced by MitoTracker staining and transmission electron microscopy (TEM). As mitochondrial structure can affect mitochondrial mass, we investigated mitochondrial content in residual cells following treatments. Quantitative PCR using primers specifically recognizing nuclear (nDNA) or mitochondrial DNA (mtDNA) revealed that DNA-damaging agents increased mtDNA content but taxanes did not (Fig. 2C, S2B). Consistent with mtDNA:nDNA ratios, quantification of the number of mitochondria in MitoTracker-stained cells revealed that DNA-damaging agents increased the number of mitochondria (Fig. 2D).

**Figure 2.**
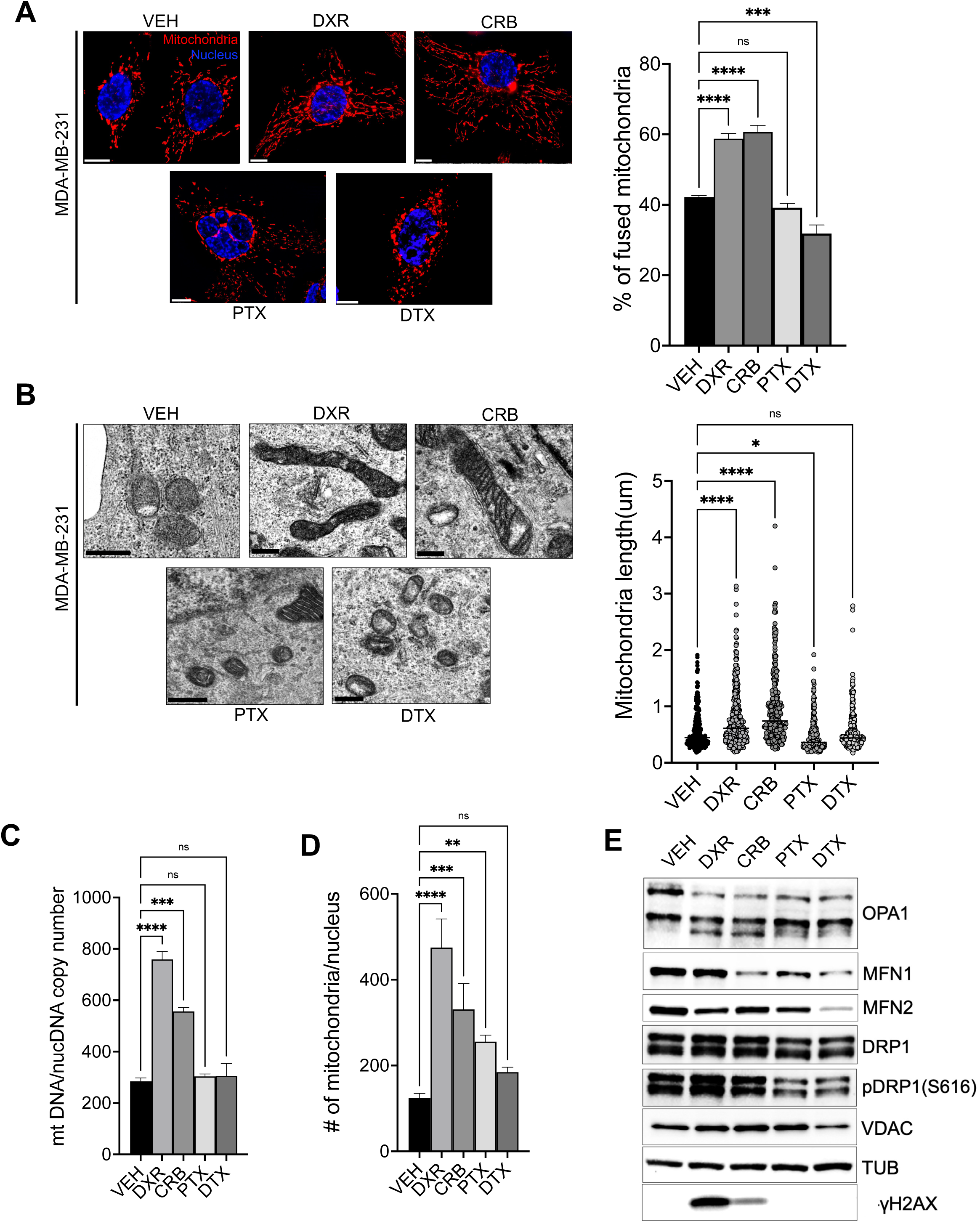
Mitochondrial structure is altered in residual TNBC cells surviving conventional chemotherapy treatments. (A) Representative fluorescence images of MDA-MB-231 cells treated with vehicle, DXR, CRB, PTX, or DTX at the IC_50_ with quantified fused mitochondrial morphology subtype using MicroP, Data are represented as the mean ± SEM (n>=3, at least 30 cells per experiment), Scale bar is 10 μm. Red fluorescence, mitochondria; blue fluorescence, DAPI-labeled nuclei. (B) Representative transmission electron micrographs of MDA-MB-231 cells (left) and quantified data (right), scale bar is 500 nm. (C) quantified mtDNA copy number: nuclear DNA copy number ratio, n=3, *\*\*\*\*P* < 0.0001, ***<0.001, **<0.01, *<0.05 by one-way ANOVA. (D) The total number of mitochondria per nucleus from MDA-MB-231 cells labeled with MitoTracker and DAPI was quantified by MicroP, *\*\*\*\*P* < 0.0001,***<0.001, **<0.01, *<0.05 by one-way ANOVA (E) Western blot analysis of mitochondrial morphology regulators. Data are representative of at least 5 independent experiments.

We then analyzed the levels of key proteins regulating mitochondrial fusion (MFN1, MFN2 and OPA1), and mitochondrial fission (DRP1 and phosphorylated DRP1) in residual cells. We observed an increase of the short isoform of OPA1 in residual MDA-MB-231 cells treated with DNA-damaging agents but not with taxanes (Fig. 2E). Further, residual SUM159pt cells following taxane treatment exhibited reduced OPA1 levels but following DNA-damaging agent treatment exhibited a slight increase in OPA1 level (Fig. S2C). OPA1 is regulated by alternative pre-mRNA splicing and proteolytic cleavages, producing isoforms with different roles in regulating mitochondrial fusion, cristae structure, and mtDNA maintenance. We observe an increase in the shortest isoform (‘isoform E’) in residual cells following DNA-damaging chemotherapy treatment. This isoform has been shown to be critical for inner mitochondrial membrane structure and mitochondrial energy production^25,28^. Other mitochondria structure regulating proteins did not exhibit clear associations with the mitochondrial structure changes we observed upon chemotherapy treatment (Fig. 2E, S2C). Together, these data indicate that distinct classes of chemotherapies have distinct impacts on mitochondrial morphology, and this is associated with altered OPA1 protein.

To address potential mechanisms underpinning the differences in mitochondrial phenotypes observed between different classes of chemotherapies, we assayed mitochondrial DNA damage by long amplification PCR of an ∼8 kb portion of the mitochondrial genome followed by qPCR to quantify long amplicons that were undamaged. We observed that DNA-damaging chemotherapies, but not taxanes, significantly induced mtDNA damage (Fig. S3A). Similarly, DNA-damaging agents, but not taxanes, induced substantial nuclear DNA damage as evidenced by γ-H2AX levels (Fig. 2E). Furthermore, we analyzed reactive oxygen species (ROS) in residual cells, revealing significant increases only following treatment with taxanes, not DNA-damaging agents (Fig. S3B). Thus, differential induction of ROS, mtDNA damage, or nDNA damage could underpin the differing effects on mitochondrial structure and OXPHOS induced by chemotherapies.

### Mitochondrial morphology impacts bioenergetics of TNBC cells

Based on the association between mitochondrial structure and metabolism we observed, we then sought to determine the functional relationship between mitochondrial structure and mitochondrial metabolism in TNBC cells. We thus examined the requirement of OPA1 and DRP1 for mitochondrial structure and bioenergetics in MDA-MB-231 cells. To ascertain the impacts of mitochondria structure alteration in viable cells that were not undergoing cell death, we treated cells with a sub-lethal dose of MYLS22, a first in class selective inhibitor of OPA1^39^ (Fig. 3A). 48 hours of MYLS22 treatment resulted in mitochondrial fragmentation as well as reduced basal and maximal OCR (Fig. 3A-B). Conversely, treatment with a sub-lethal dose of Mdivi-1 (Fig. 3A), a small-molecule inhibitor of the mitochondrial fission protein DRP1^40^, resulted in mitochondrial elongation and heightened basal and maximal OCR (Fig. 3A-B). We then conducted stable isotope metabolomic flux tracing in ^13^C_6_ -glucose or ^13^C_5_ -glutamine in SUM159pt cells treated with a sub-lethal dose of MYLS22 or Mdivi-1. We found substantial reduction of glucose-driven, but not glutamine-driven, TCA cycle flux in MYLS22-treated cells (Fig. 3C). Mdivi-1 did not significantly alter glycolytic nor TCA cycle flux, suggesting Mdivi1-induced mitochondrial fusion was not sufficient to drive these metabolic pathways. These findings suggest that OPA1 function may be required for glucose flux into the TCA cycle and resulting OXPHOS.

**Figure 3.**
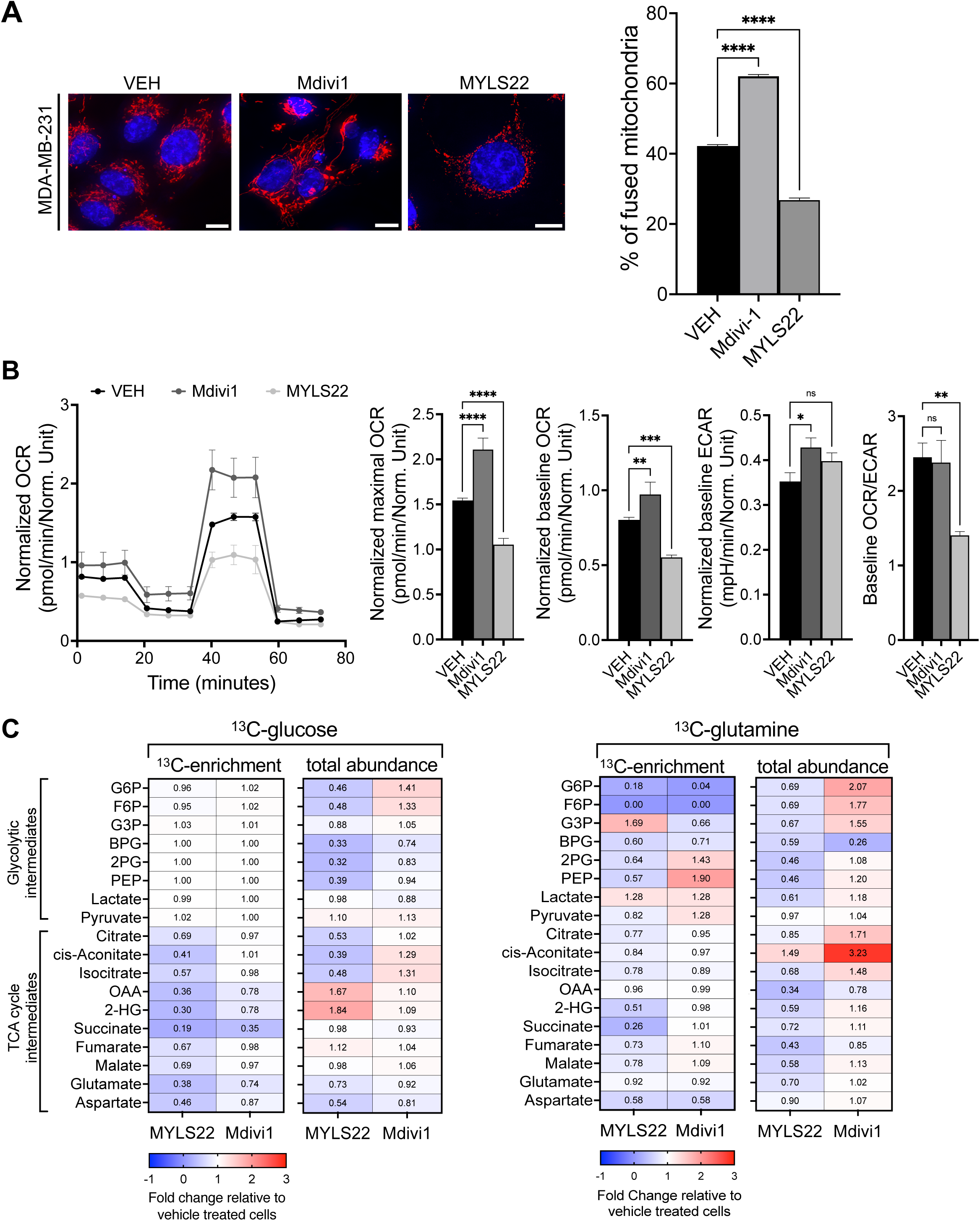
Pharmacologic perturbation of mitochondrial morphology functionally impacts mitochondrial metabolism in TNBC cells. (A) Representative fluorescence images of the cells treated with vehicle, Mdivi-1, or MYLS22 then stained with MitoTracker and DAPI, with quantified mitochondria morphology subtypes from MicroP analysis, *n* >30 cells. Scale bar is 10 μm. Red fluorescence, mitochondria; blue fluorescence, DAPI-labeled nucleus. (B) Representative Seahorse Mito Stress assay showing OCR, maximal OCR, basal OCR, basal ECAR and OCR/ECAR ratio in MDA-MB-231 cells treated vehicle, Mdivi-1, or MYLS22. \**\*\*\*P* < 0.0001, ***<0.001, **<0.01 by one-way ANOVA test. Data are representative of at least 3 independent experiments. (C) ^13^C-glucose (left) derived or ^13^C-glutamine (right) derived-enrichment or total abundance of glycolytic and TCA cycle intermediates in SUM159pt cells treated vehicle, MYLS22, or Mdivi-1. Data were normalized to labeling of vehicle treated cells.

We complemented these pharmacologic studies by genetically perturbing levels of mitochondrial fission and fusion proteins in MDA-MB-231 cells. siRNA knock-down (KD) of OPA1 (Fig. 4A, upper panels) resulted in mitochondrial fragmentation and reduced OCR (Fig. 4B-C). Conversely, siRNA KD of DRP1 (Fig. 4A, lower panels) resulted in mitochondrial elongation and increased OCR (Fig. 4B, D). Our findings reveal that in TNBC cells, mitochondrial fusion can promote OXPHOS, whereas fragmented mitochondria have reduced OXPHOS.

**Figure 4.**
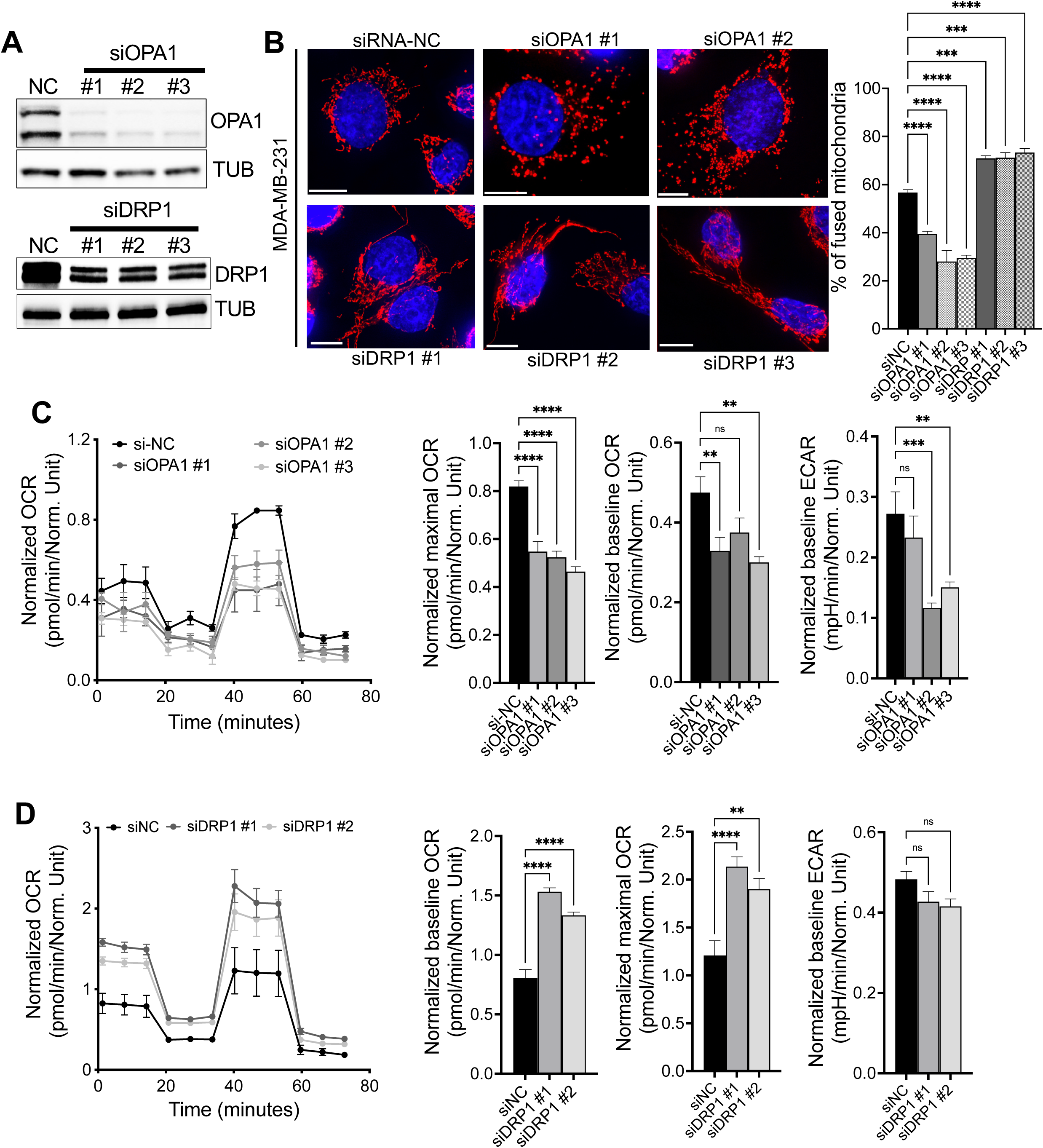
Genetic perturbation of mitochondrial morphology dictates mitochondrial metabolism in TNBC cells. (A) Immunoblotting verifying knockdown of *OPA1* (siOPA1, top panel) or *DRP1* (siDRP1, bottom panel) in MDA-MB-231 cells. (B) Representative fluorescence images of the cells transfected with NC (non-targeting siRNA), siOPA1 or siDRP1 with MitoTracker and DAPI staining, with quantified mitochondria morphology subtypes, Data are represented as the mean ± SEM (n>=3, at least 30 cells per experiment), \**\*\*\*P* < 0.0001, ***<0.001, **<0.01 by one-way ANOVA test. Scale bar is 10 μm. Red fluorescence, mitochondria; blue fluorescence, DAPI-labeled nucleus. (C-D) Representative Seahorse Mito Stress assay showing OCR, maximal OCR, basal OCR, basal ECAR in siOPA1 or siDRP1 cells compared with controls. \**\*\*\*P* < 0.0001, ***<0.001, **<0.01 by one-way ANOVA test. Data are representative of at least 3 independent experiments.

### Mitochondrial morphology impacts chemo-sensitivity of TNBC cells

We sought to determine the requirement of mitochondrial fusion for DNA-damaging chemotherapy resistance. KD of OPA1 sensitized MDA-MB-231 cells to doxorubicin treatment without substantially impacting cell growth in the absence of chemotherapy treatment (Fig. 5A-B). We then tested the requirement for mitochondrial fusion and fission in mediating survival of residual TNBC cells that are not killed by conventional chemotherapy treatment. We first treated MDA-MB-231 with an IC_50_ dose of doxorubicin for 48 hours followed by a sub-lethal dose of the selective OPA1 inhibitor MYLS22. MYLS22 significantly increased doxorubicin-induced cell killing (Fig. 5C) and reduced doxorubicin-mediated mitochondrial elongation and OXPHOS (Fig. 5D-E). Of note, we observed that order of addition of agents impacting mitochondria is a crucial consideration. Specifically, treatment of cells in specific order with MYLS22, inducing fission, followed by doxorubicin, inducing fusion, enhanced cell growth compared to cells treated with doxorubicin alone (Fig. S4B). Treatment of cells with doxorubicin followed by MYLS22 or given concurrently with MYLS22 drastically decreased cell growth (Fig. S4A). Interestingly, cells treated with doxorubicin, inducing fusion, followed by Mdivi-1, further inducing fusion, were less sensitive to doxorubicin than were vehicle-treated cells (Fig. S4B). Together, these results indicate pharmacologic perturbation of mitochondrial structure can overcome chemotherapy-induced adaptations in TNBC cells when administered rationally based on our understanding of chemotherapy-induced mitochondrial adaptations.

**Figure 5.**
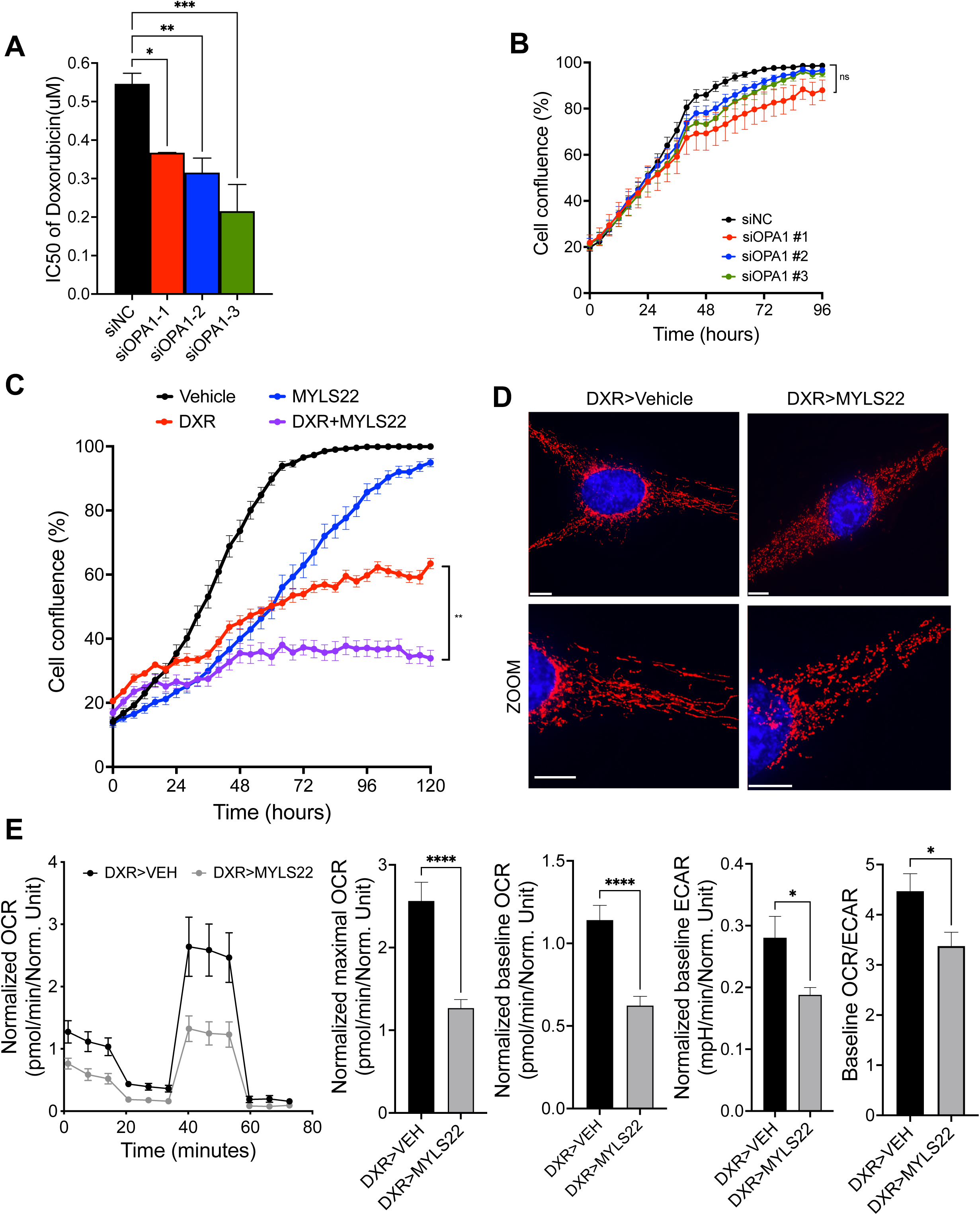
Mitochondrial morphology impacts chemo-sensitivity of TNBC cells. (A) The IC_50_ of DXR in MDA-MB-231 cells transfected with three different siOPA1 variants compared to control (NC, non-targeting siRNA) was measured by Cell Titer Glo luminescence. Data are represented as the mean ± SEM, n=3, \**\*\*P* < 0.001, **<0.01, *<0.05 by one-way ANOVA test. (B) IncuCyte time lapse imaging of confluence of MDA-MB-231 transfected with three different siOPA1 variants compared to control (NC) (C) IncuCyte analysis of MDA-MB-231 treated with vehicle, MYLS22 only, DXR only, or DXR followed by MYLS22 treatment at 48 hours. (D) Representative fluorescence images of the cells treated with DXR followed by vehicle or MYLS22 with MitoTracker and DAPI staining, Scale bar is 10 μm. Red fluorescence, mitochondria; blue fluorescence, DAPI-labeled nucleus. (E) Representative Seahorse Mito Stress assay showing OCR, maximal OCR, basal OCR, basal ECAR and OCR/ECAR ratio in MDA-MB-231 cells treated with DXR followed by vehicle or MYLS22, \**\*\*\*P* < 0.0001, *<0.05 by one-way ANOVA test. Data are representative of at least 3 independent experiments.

### Mitochondrial adaptations in a PDX model of post-chemotherapy residual TNBC

The standard front-line DNA damaging combination chemotherapy regimen AC remains a key part of TNBC neoadjuvant treatment^41^. We previously developed and deeply characterized PIM001-P, an orthotopic PDX model of primary treatment-naïve TNBC^7^. Treatment of this model with AC induced a residual tumor drug-tolerant state with heightened susceptibility to inhibition of OXPHOS using the ETC Complex I inhibitor IACS-010759^7,8^. To analyze mitochondria in this PDX model, we collected tumors from NOD/SCID mice bearing orthotopic PIM001-P tumors treated without (pre-treated) or with AC following two days (ACd2), 21 days (residual tumors), and 50 days (regrown tumors) (Fig. 6A). We then directly measured mitochondrial OXPHOS capacity in purified tumor cells from freshly dissociated pooled tumors from each timepoint that had been depleted of mouse stroma as we previously described^7^. Seahorse analysis revealed residual tumor cells had heightened basal and maximal OCR (Fig. 6B). Immuno-histochemical (IHC) staining of tumor sections with an antibody specifically recognizing human mitochondria, thus excluding mouse stroma, revealed increased mitochondrial content specifically in residual tumor cells by machine learning-based quantification (Vectra, Akoya; Fig. 6C-D). We corroborated this by qPCR of mtDNA extracted from purified human tumor cells (Fig. 6E). Western blot analysis of purified tumor cells revealed increased OPA1 in residual tumor cells (Fig. 6H). Further, TEM analysis of these tumors revealed increased mitochondrial length in residual tumor cells (Fig. 6F-G). Co-immunofluorescence of VDAC to mark mitochondria and OPA1 revealed increased co-staining in residual tumors (Fig. 6I and S5). Together, these findings suggest that increased OXPHOS is accompanied by increased mitochondrial fusion, mitochondrial content, and OPA1 levels in an *in vivo* PDX model of residual TNBC.

**Figure 6.**
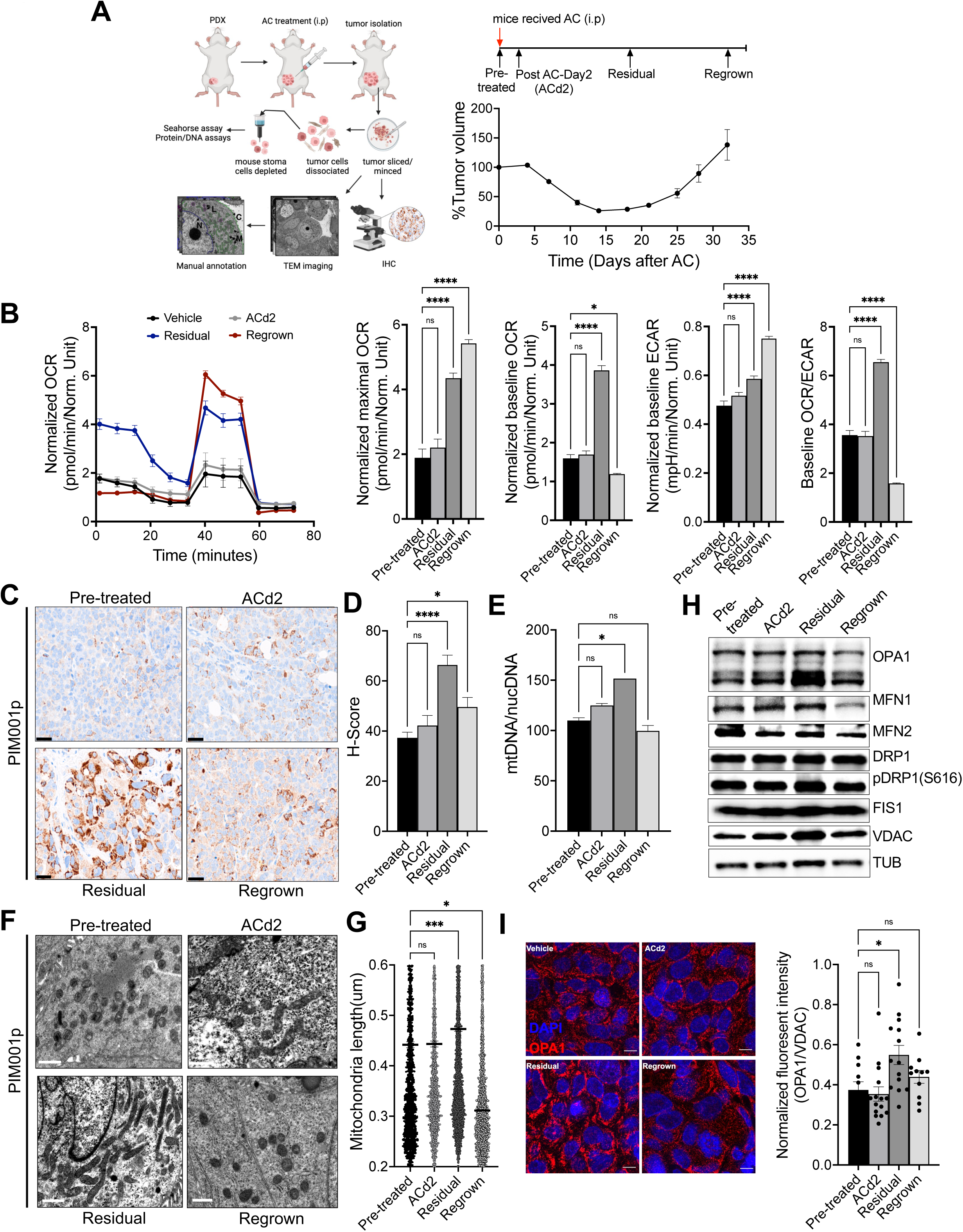
Mitochondrial adaptations in a PDX model of post-AC residual TNBC. (A) Schematic diagram of the experimental approach for tumor isolations and experimental procedures performed on PDX tumors isolated from NOD/SCID mice bearing PIM001-P tumors treated with a single AC dose (day 0; 50 mg/kg C + 0.5 mg/kg A). (B) Seahorse Mito Stress assay of the purified human tumor cells from pre-treated, post-AC treated for 2days (ACd2), residual (21 days), and regrown (35 days) PIM001p tumors. n=3, (C) Representative IHC images of PIM001-P tumors stained with human-specific mitochondria antibody and (D) quantified by Vectra 3 imaging. Scale bar is 20μm. (E) Quantified mtDNA:nDNA ratio of DNA prepared from snap-frozen purified tumor cells. n=3-4 tumors were pooled per time point. (F) Representative TEM images of PIM001-P Pre-treated(vehicle), AC-treated residual and AC-treated regrown tumors. Scale bar is 1 mm. (G) The average mitochondria length was calculated from TEM images. Approximately 3000 mitochondria per group were analyzed across 3 biological replicate mice per group. (H) Western blotting was conducted on protein lysates made from 3-4 pooled tumor from each timepoint. (I) Representative immunofluorescence images of the tumors stained with OPA1 antibody (red) and nucleus dye DAPI (blue), scale bar is 10 μm. Quantified fluorescent intensity of OPA1 normalized to VDAC (mitochondria marker) is shown in a bar graph. \**\*\*\*P* < 0.0001, ***<0.001, **<0.01, *<0.05 by one-way ANOVA test (n=3)

### Inhibition of OPA1 perturbs residual tumor regrowth in a PDX model of residual TNBC

Based on our data revealing mitochondrial structure dynamics accompany the residual tumor phenotype in the PDX PIM001-P, we tested the hypothesis that residual tumors would be uniquely susceptible to perturbation of mitochondrial fusion. We conducted a preclinical residual tumor trial to target mitochondrial fusion modeled after our prior study with IACS-010759^7^. We orthotopically engrafted NOD/SCID mice with PIM001-P tumors. Treatment of non-chemotherapy-treated mice with the OPA1 inhibitor MYLS22 as a single agent had no effect on tumor growth (Fig. 7A,C). We treated a subset of mice with AC to induce the ‘residual tumor state’ as previously described^7^, then began MYLS22 treatment of residual tumor bearing mice 21 days after AC treatment. We found that the regrowth of AC-treated residual tumors was significantly delayed by MYLS22 (Fig. 7B,C). In all groups, mice body weight was unaffected (Fig. S5B). These findings reveal that residual PIM001-P tumors had an enhanced dependency on mitochondrial fusion compared to treatment-naïve tumors.

**Figure 7.**
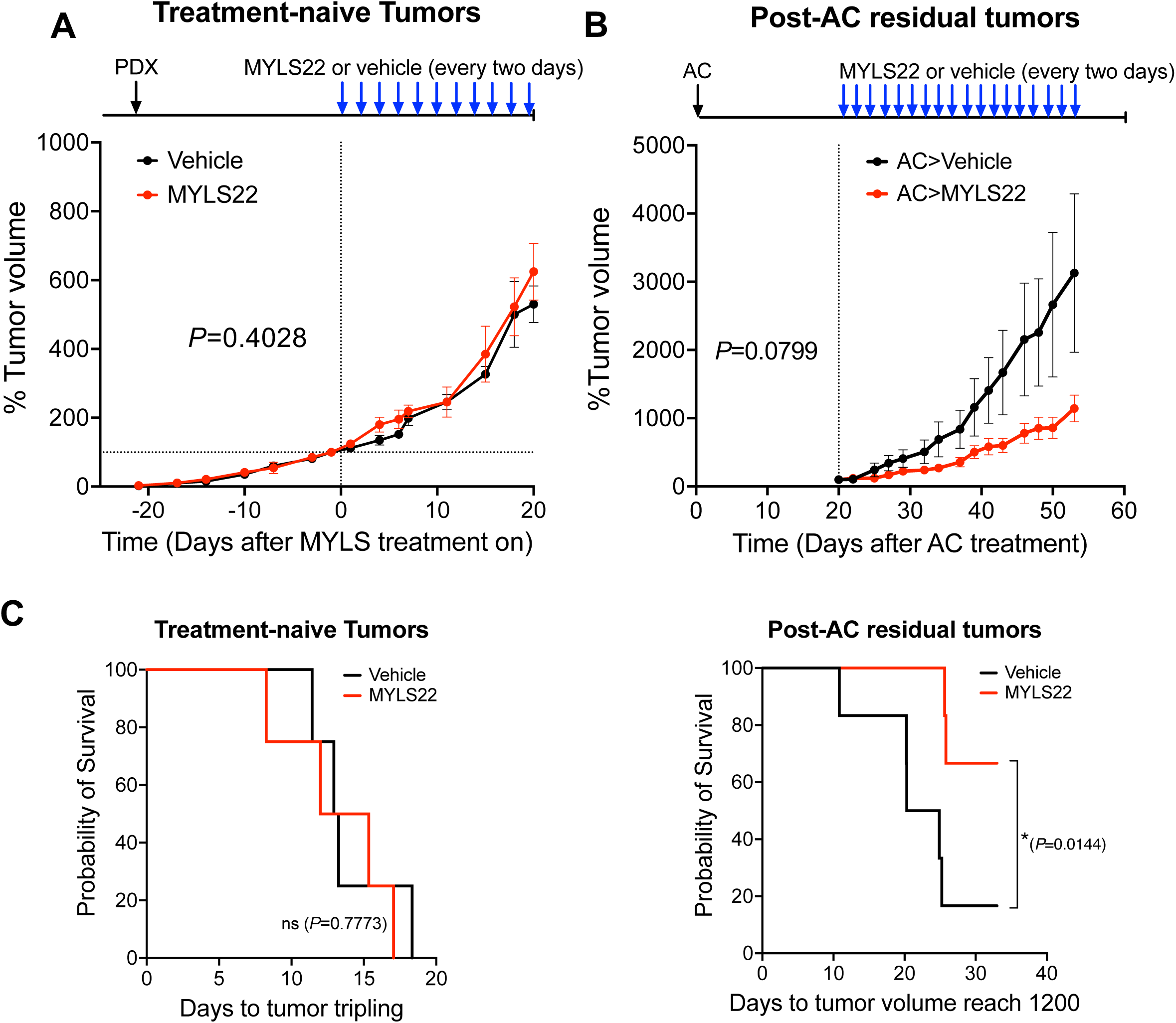
Inhibition of OPA1 perturbs residual tumor regrowth in a PDX model of residual TNBC. (A-B) normalized tumor growth curve of NOD/SCID mice bearing PIM001-P tumors that were treated with an inhibitor of OPA1, MYLS22, or vehicle in the treatment-naïve (A) or in the residual setting after AC treatment (B). Each day of MYLS22 treatment is indicated by the blue arrow. Error bars represent SEM (n=4–6 per group, *t-*test). Top plots display tumor growth, *P*-value calculated from *t-*test. (C) Survival plots were analyzed by Wilcoxon test.

## DISCUSSION

While some tumors exhibit high rates of glycolysis, mitochondrial OXPHOS is critical for tumor progression and therapy resistance in breast cancer^7,9^, melanoma^42^, pancreatic cancer^43^, and leukemia^11^. In TNBC, ∼45% of patients will harbor substantial residual tumor burden following treatment with NACT. At present, there is no accepted replacement for standard chemotherapy in TNBC. Thus, it is crucial we delineate mechanisms driving residual tumor cell survival that may ultimately present us with therapeutic targeting opportunities. TNBCs can rely on OXPHOS for survival^7,9,44^, driven by energy sources such as glucose or fatty acids^45,46^. In fact, mitochondrial transplantation experiments demonstrated these organelles are both necessary and sufficient to confer aggressive phenotypes of TNBC cells^45^. Metabolic subtyping of 465 patient biopsies revealed OXPHOS is the most upregulated metabolic pathway in TNBC tumors compared to normal mammary tissue^47^ and this was recently corroborated by transcriptomics^48^. Thus, finding ways to target OXPHOS dependency holds great promise to impact TNBC outcomes.

AC remains a key front-line combination chemotherapy for the neoadjuvant management of TNBC^41^. Therefore, understanding mechanisms by which tumor cells survive AC treatment may open new therapeutic avenues. In our prior work we modeled residual disease by treating orthotopic PDX models derived from treatment naïve TNBC patients with AC. We found residual tumors in PDX models underwent a switch from glycolysis to OXPHOS. Despite the limitation that PDX models lack a fully intact immune system, we were able to corroborate this finding using gene expression data^49^ from serial TNBC patient biopsies. We conducted ‘preclinical trials’ in PDX models modeled after a human neoadjuvant clinical trial^50^ and demonstrated OXPHOS is a therapeutic vulnerability in the residual setting of TNBC^7^ by treating with an inhibitor of ETC Complex I, IACS-010759^8^. Together, these findings led us to investigate the mechanistic underpinnings of this metabolic dependency. In the present study, we found the heightened basal and maximal OXPHOS rate is associated with mitochondrial elongation in residual tumors of this PDX model. Based on our observation residual tumors had increased OPA1, we conducted a preclinical trial with MYLS22, a first in class specific inhibitor of OPA1^39^, revealingnMYLS22 had greater efficacy in the residual setting than it did in the naïve setting. These results suggest targeting mitochondrial structure adaptations that enable pro-survival metabolic rewiring may be a promising avenue to kill residual TNBC cells.

OPA1 orchestrates mitochondrial genome maintenance, cristae structure, ETC complex assembly, and mitochondrial inner membrane fusion^25,26^. Skipping of alternative exon 5b followed by proteolytic cleavage by OMA1 produces the short ‘E’ protein isoform, required for proper inner mitochondrial membrane structure and ETC complex formation^25,28^. In some stress conditions, OMA1 activity is enhanced, resulting in accumulation of short OPA1 isoforms^29^. Our *in vitro* data provide evidence that OPA1 short isoform production is increased in residual tumor cells following DNA-damaging chemotherapy treatments and is correlated with mitochondrial elongation and increased OXPHOS. The complex regulation of OPA1 alternative splicing and proteolytic cleavage are poorly understood and have not yet been investigated in TNBC. A greater understanding of these mechanisms is expected to provide further therapeutic targeting opportunities for chemoresistant TNBC.

Our metabolomic flux tracing experiments revealed that DNA-damaging and taxane chemotherapies increased TCA cycle flux. However, this increased TCA cycling only manifest as increased OXPHOS and mitochondrial fusion in DNA-damaging chemotherapy-treated cells but not taxane treated cells. Thus, we hypothesize that mitochondrial morphological status plays a critical role in optimizing OXPHOS activity downstream of the TCA cycle. Indeed, flux analysis of cells treated with the fission inhibitor (thus increasing fusion) Mdivi-1 revealed no change in TCA cycle labeling. On the other hand, the fusion inhibitor MYLS22 caused decreased labeling of TCA cycle intermediates. These findings suggest that mitochondrial fusion is necessary, but not sufficient, for increasing OXPHOS downstream of increased TCA cycling.

The mechanistic basis for the differential mitochondrial effects caused by DNA-damaging (carboplatin and doxorubicin) versus taxane (paclitaxel and docetaxel) chemotherapies is yet unclear. DNA-damaging agents induced both nuclear and mitochondrial DNA damage, which could provide a pro-fusion and pro-OXPHOS adaptive signal through not yet understood mechanisms. Furthermore, mitochondrial dynamics are tightly linked with the cell cycle. Thus, cell cycle arrest at different phases induced by distinct classes of chemotherapies may also underly the differential mitochondrial phenotypes of residual cells. The ROS we observed specifically induced by taxanes may also be responsible for their induction of mitochondrial fission.

Perturbation of mitochondrial structure is being developed as an anti-cancer strategy. Studies have demonstrated targeting mitochondrial fission or fusion can perturb tumor growth without substantial host toxicity in preclinical models^33,36,51^. In the clinic, the MFN2-inducing drug Leflunomide^52^ is approved for treatment of arthritis and second mitochondrial-derived activator of caspases (SMAC) mimetics such as Birinapant, shown to be toxic in tumor cells with defective mitochondrial structure^51^, are in trials for various tumor types^53^. Silibinin, a natural flavonolignan that inhibits mitochondrial fusion through poorly understood mechanisms, is safe and well tolerated in human patients^54^. While none of these current clinical agents are selective inhibitors of mitochondrial structure dynamics, we hypothesize there is a safe therapeutic window in residual TNBCs for targeting mitochondrial structure. These data should support continued development of specific inhibitors of mitochondrial fusion for clinical use.

## MATERIALS AND METHODS

### Cell culture

All cell lines were purchased directly from ATCC (American Type Culture Collection) except for SUM159pt which was purchased from BioIVT. MDA-MB-231 cells were cultured in RPMI-1640 (Gibco, 11879020) supplemented with 5 mM glucose (Gibco, A2494001), 10% fetal bovine serum (R&D, S11550), and 1x antibiotic-antimycotic solution (Corning, 30-004-CI). SUM159pt cells were cultured in RPMI-1640 (Gibco, 11879020) supplemented with 5 mM glucose (Gibco, A2494001), 5ug/ml Insulin (Sigma, I9278), 1ug/ml hydrocortisone (Sigma, H4001), 5% fetal bovine serum (R&D, S11550), and 1x antibiotic-antimycotic solution (Corning, 30-004-CI). All the cells were incubated at 37°C with 5% CO_2_ and 95% relative humidity.

### Gene silencing

Cells were seeded and maintained in growth medium without antibiotics prior to siRNA transfection. Cells were transiently transfected with 10nM siRNA against human *OPA1* (5’-AAGAAUUUUCCCGCUUUAUGA-3’, 5’-GACAUAUUUGAUAAACUUAAA-3’, 5’-GUUAUCAGUCUGAGCCAGG-3’), human *DRP1* (5’-GCAGAAGAAUGGGGUAAAU-3’, 5’-UCCGUGAUGAGUAUGCUUU-3’, 5’-GAGGUUAUUGAACGACUCA-3’) non-targeting scrambled siRNA (Sigma, SIC001-1NMOL) for 48-72 hours using Lipofectamine RNAiMAX (Invitrogen, 13778150) in OPTI-MEM media (Gibco, 11058021) according to the manufacturer’s instructions. To knock-down OPA1 gene expression, MDA-MB-231 cells were transduced with GFP labelled shRNA lentiviral particles (pRSIT17-U6Tet-sh-CMV-TetRep-2A-TagGFP2-2A-Puro, 1.30 × 10^7^ TU, customized by Cellecta (Mountain View, CA) against OPA1 (5’-GTTATCAGTCTGAGCCA-3’) or non-targeting scramble sequence (5’-CAACAAGATGAAGAGCACCAA. ∼2.0×10^6^ MDA-MB-231 cells were seeded with growth media without antibiotics per well in a 6-well plate and incubated at 37°C overnight. On the following day, cells were treated with 5ug/ml polybrene (Sigma, TR-1003-G) and shRNA with (1-5) MOI (Multiplicity of Infection). After 48-72hrs, puromycin (2ug/ml, Sigma, A1113802) antibiotic selection for identifying resistant colonies was initiated. Non-transduction cells were used to verify puromycin selection. To induce *OPA1* knockdown, cells were subsequently maintained in growth medium supplemented with doxycycline (1ug/ml, Sigma, D9891). Transduction efficiency was monitored by IncuCyte ZOOM (Essen BioScience, Ann Arbor, MI)-based GFP fluorescence imaging.

### *In vitro* drug treatments

With the exception of carboplatin (solutions were freshly prepared from powder), we prepared stock solutions of 10 mM in DMSO and further dilutions were prepared in proper growth medium. Cells were seeded at ∼20% confluence and maintained for 48hrs prior to treatment. Cells were then treated with: doxorubicin (final conc. at 50-100nM, Sigma, 44583, protected from light), carboplatin (final conc. at 50-100uM, Selleck Chemicals, Houston TX, S1215), paclitaxel (final conc. at 1-10nM, Selleck Chemicals, Houston TX, S1150), docetaxel (final conc. at 1-5nM, Selleck Chemicals, Houston TX, S1148), Mdivi-1 (final conc. 1-5μM, Sigma, M0199), or MYLS22 (final conc. 50-100μM, MedChem Express, Monmouth Junction, NJ, HY-136446) for 24-72 hours prior to downstream analyses.

### Animal Studies

This study was carried out in accordance with the *Guide for the Care and Use of Laboratory Animals* from the National Institutes of Health (NIH) IACUC. The protocol was approved by the IACUC at BCM (protocol AN-8243). Mice were euthanized when they reached defined study or ethical end points. Euthanasia was conducted as recommended by the Association for Assessment and Accreditation of Laboratory Animal Care International.

PDX models were originally obtained from the University of Texas MD Anderson Cancer Center, where they were generated and characterized, through a materials transfer agreement. For this study, PIM001-P was propagated as previously described^1^. Briefly, cryo-preserved PDX cell suspensions were quickly thawed, then washed with Dulbecco’s modified Eagle’s medium (DMEM):F12 (Cytiva HyClone, SH30023.01) supplemented with 5% FBS. Viable cells were counted by staining with AOPI dye (Nexcelom Bioscience, CS2-0106) on a Cellometer K2 (Nexcelom Bioscience). For injection into mammary glands, 0.5-1.0 million viable tumor cells were suspended in a total of 20 μl (1:1 volume mixture of medium and Matrigel (Corning, 354234). Suspensions were then immediately injected unilaterally into the fourth mammary fat pads of 5- to 8-week-old NOD/SCID mice [NOD.CB17-Prkdc^scid^/NcrCrl, Charles River, National Cancer Institute (NCI) Colony].

Chemotherapy treatment of PDX models was conducted as previously described. Briefly, Adriamycin (doxorubicin, ChemieTek, CT-DOXO) was solubilized in sterile water for injection immediately prior to administration at 0.5 mg/kg. Cyclophosphamide (Sandoz, 0781-3244-94) was solubilized in sterile water for injection immediately prior to administration at 50 mg/kg. Each solution was administered by intraperitoneal injection separately at a dose volume of 5 to 10 ml/kg.

MYLS22 was solubilized with corn oil by sonication in a 37°C water bath to make a 200 mM stock solution. The stock solution was stored at -80°C and used within 5 days after preparation. The fourth mammary fat pads of NOD/SCID mice were unilaterally engrafted with 0.5 × 10^6^ PIM001-P tumor cells per mouse. When tumors reached an average of 150mm3, mice were randomized to four treatment groups based on tumor volume: 4.5 ml/kg vehicle (Corn Oil, Sigma, C8267-500ML) peritumoral injection every other day, 10 mg/kg MYLS22 peritumoral injection at 4.5 ml/kg every other day until end of the experiment, a single dose of AC as described above, or a single dose of AC followed by MYLS22 treatment of residual tumors beginning 21 days after AC treatment. Residual tumors received MYLS22 every two days until end of the experiment

### Dissociation of PDX tumor cells

Tumors were minced using scalpels and digested in Dulbecco’s modified Eagle’s medium (DMEM):F12 (Cytiva HyClone, SH30023.01) supplemented with 5% FBS, 0.45% collagenase A (Roche, 1088793), 0.086% Hyaluronidase (Sigma, H3506) and 2% BSA (Sigma, A9418) for 4-6 hours at 37°C with rotation. Cells were then collected by centrifugation at 170 x *g* for 8minutes at room temperature. Cells were resuspended in red blood cell (RBC) lysis buffer (Sigma, R7757), incubated for 3 minutes with periodic inverting and added the equal volume of epithelial cell growth medium (Dulbecco’s modified Eagle’s medium (DMEM):F12 (Cytiva HyClone, SH30023.01) supplemented with 5% FBS and 1x antibiotics). Cells were collected by centrifugation and treated with trypsin for 3minutes. Trypsin was inactivated by adding equal volume of epithelial cell growth medium, centrifugate cells at 170 x *g* for 5minutes and then discard supernatant. The cells were treated by Dispase solution (Stemcell Technologies, NC9886504) with DNase I (Stemcell Technologies, NC9007308) for 5 minutes. Dispase was inactivated by adding equal volume of growth medium. The cell suspension was applied to a cell strainer (70 μm and then 40 μm, Corning) and centrifuged at 1000rpm for 5 minutes. After determining total cell counts, mouse stromal cells were depleted using mouse cell depletion kit (Miltenyi Biotec 130-104-694) according to manufacturer’s instructions. The purified tumor cells were resuspended in growth medium and used for further analyses.

### Validation of cell lines and PDX models

Cell cultures were tested for mycoplasma contamination each quarter by PCR (the Universal Mycoplasma detection kit, ATCC, 30-1012K) and were confirmed mycoplasma-free throughout the duration of this study. Short-tandem repeat (STR) DNA fingerprinting was performed on DNA extracted from cell lines and on PDX model tumors the Cytogenetics and Cell Authentication Core (CCAC) at M.D. Anderson Cancer Center. The Promega 16 High Sensitivity STR Kit (Catalog # DC2100) was used for fingerprinting analysis, and profiles were compared to online search databases (DSMZ/ ATCC/ JCRB/ RIKEN).

### Transmission electron microscopy (TEM) and analysis

Cell suspensions were centrifugated at 800 x *g* at room temperature for 10 minutes to make a pellet. The supernatant was removed and replaced with ∼1ml of Trump’s fixation solution (2% paraformaldehyde, 3% glutaraldehyde in phosphate buffer). The cells were gently re-suspended, transferred to new Eppendorf tube, and allowed to fix at room temperature with gentle rocking. For tumor tissues, immediately following animal euthanasia, mammary tumors were resected from mice and a ∼1 mm-tick tumor slice was placed in Trump’s fixation solution and fixed for 24 hours at room temperature with rocking. The necrotic areas that usually found in the core region of solid tumors were avoided. The initial fixed samples then were post-fixed in 1% buffered osmium tetroxide, dehydrated in graded ethanol, and embedded in Polybed 812 resin. Ultra-thin sections were obtained using a Leica UC 7 and sections were stained in 1% aqueous uranyl acetate and Reynold’s lead citrate. Images were obtained using a FEI Spirit Tecnai transmission electron microscope equipped with and Eagle camera and FEI Tia image acquisition software. Total ∼65 images taken at magnifications (400-5000x) were taken per tumor sample. Images were annotated and then quantified by QuPath software^55^.

### Fluorescence microscopy imaging and analysis

For fluorescence analysis of mitochondrial morphology, a mitochondria-specific fluorescent dye MitoTracker CMXRos (Thermo Fisher Scientific, M7514) was used to monitor mitochondrial morphology according to the manufacturer’s instructions. Briefly, cells were then seeded on coverslips coated with Poly-D-lysine (Sigma, P6407) and treated with fresh growth media containing 100ng/ml MitoTracker CMXRos (Invitrogen, M7512) at 37°C for 15-30 minutes. Cells then were washed twice with growth media and fixed with pre-warmed media containing 3.7% formaldehyde solution for 15 minutes at 37°C. After fixation, cells were washed with PBS three times, treated with 0.2% of Triton X-100 for 10 minutes at room temperature, then treated with 300nM of DAPI in PBS (Invitrogen, D3571). Coverslips were mounted on glass slides with 25μl of Fluoromount-G™ Mounting Medium (Invitrogen, 00-4958-02). Fluorescence imaging was carried out on DeltaVision high resolution fluorescent microscope (GE Healthcare) equipped with SoftWork software. Raw images were processed using ImageJ. Mitochondria shape and number were analyzed by the MAT-LAB based macro, MicroP according to the author’s instructions^56^. Up to 10 images at 60x magnification per sample were analyzed.

### DNA extraction and mtDNA quantification

Total DNA was extracted using the Mag-Bind® Blood & Tissue DNA HDQ 96 Kit according to manufacturer’s instructions with some slight modifications. Purified DNA samples were quantified by NanoDrop 2000 (Thermo Scientific). Quantitative PCR was performed with the Universal SYBR Green Supermix (Bio-Rad, 1725121), 2 to 4 ng of DNA, and 1μl of primers recognizing mitochondrial ND1 (forward: 5’-ATGGCCAACCTCCTACTCCT-3’, reverse: 5’-TAGATGTGGCGGGTTTTAGG-3’), mitochondrial ND6 (forward: 5’-TGGGGTTAGCGATGGAGGTAGG, reverse: 5’-AATAGGATCCTCCCGAATCAAC-3’), nuclear RGPD1 (forward 5’-GTGGAGCCACTGAGAATGGT-3’, reverse: 5’-GCATGCCTGGCTGATTTTAT-3’), or nuclear FUNDC2P2 (forward 5’-TGAGTCAGTGGACCTTGCAG-3’, reverse: 5’-CAGAATGGTTTGCAAGCTGA-3’). We used primers to compare the levels of two nuclear genes (RGPD1 and FUNDC2P2) to two mitochondrial genes (ND1 and ND6) in a real-time quantitative polymerase chain reaction. Mixtures of 2 to 4 ng of DNA, primers and 2xSYBR master mix (Bio-Rad, 1725121) were analyzed with the cycling conditions: 95 °C for 10 minutes followed by 40 Cycles of: 95 °C for 15 seconds 64 °C for 1 minute. Relative copy number was calculated using the relative 2^-ΔΔCt^ method.

### Mitochondrial DNA damage assay

Long amplification PCR (LA-PCR) was performed in a 25 μl reaction on mixture containing 2.5units LongAmp Taq DNA Polymerase, 1x LongAmp Taq Reaction Buffer, 300 μM of dNTPs (New England Biolabs, E5200S), 0.5 μM of each primer (8kb mitochondrial DNA fragment, forward: 5’-TCTAAGCCTCCTTATTCGAGCCGA-3’, reverse: 5’-TTTCATCATGCGGAGATGTTGGATGG-3’), and 50ng of DNA template. Amplification comprised an initial denaturation at 94°C for 30 seconds, followed by 30 cycles of 10 seconds at 94°C, 15 seconds at 68°C, and 4 minutes at 72°C and then final extension at 72°C for 10minutes (MyCycler, Bio-rad). After the reaction, the PCR product was 1:10 diluted for further Real-time PCR. Real-time PCR amplifications were performed in 10μl reaction volumes containing 2μl of DNA template, 0.5uM of each primer (human MT-ND6 forward: 5’-TGGGGTTAGCGATGGAGGTAGG, reverseerse:5’-AATAGGATCCTCCCGAATCAAC-3’), 1x iTaq™ Universal SYBR® Green Supermix SYBR Universal Master Mix (Bio-rad). The following cycling parameters were used: 2 minutes at 50°C, 10 minutes at 95°C, then 40 cycles of 15 seconds at 95°C and 60 seconds at 60°C using CFX96 Touch Real-Time PCR Detection System (Bio-rad). A standard curve was derived from the serial dilutions of DNA template following concentrations: 2, 0,2, 0.02, 0.002, 0.0002, and 0.00002ng. The “threshold cycle” (Ct) values are then plotted versus the logarithm of the dilution. The long-amplified DNA content is calculated by using linear regression. If the DNA content is high, mitochondria DNA has relatively less damage.

### Western blotting

Total protein lysates were extracted with RIPA buffer (Thermo Scientific, 89901) supplemented with protease inhibitor cocktail (Roche, 11836153001) and PhosSTOP™ (Roche, 4906837001) according to the manufacturer’s guidelines. Protein lysates were quantified by Pierce BCA assay (Thermo Scientific, 23222). Protein samples were diluted in 4x sample buffer (Bio-rad, 1610747) with 10% beta-mercaptoethanol (Sigma, M6250) and denatured by heating at 95°C for 10 minutes. Diluted protein samples (15ug of protein per lane) were separated on pre-cast 4-20% gradient (Bio-Rad, 5671095) or 7.5% SDS-polyacrylamide gels (Bio-rad, 5671024) then transferred to a nitrocellulose membrane (Bio-rad, 1704271) using the Trans-Blot Turbo system (Bio-rad). After blocking with EveryBlot Blocking Buffer (Bio-rad, 12010020) for 10 minutes at room temperature, membranes were incubated with primary antibodies overnight at 4°C on a rocker, then incubated with HRP-conjugated secondary antibody for 1 hour at room temperature at a dilution of 1:10,000. Blots were washed and then detected using chemiluminescent HRP substrate detection kit (Thermo Scientific Pierce, 34076). Images were acquired on a ChemiDoc MP apparatus (Bio-rad) and ImageLab v.5.2(Bio-rad).

### Immunohistochemistry and immunofluorescent staining and analysis

PDX tumor samples were formalin fixed and processed into paraffin blocks as previously described^7^. 3μm sections were cut for histologic analyses. Antigen retrieval was performed using 1X reveal decloaker solution (Biocare Medical, RV1000M) at 98°C for 15 minutes. For co-immunofluorescence (Co-IF), rabbit anti-VDAC (Abcam, ab154856, 1:100) and mouse anti-OPA1 (BD Transduction Laboratories, 612607, 1:2000) were diluted in antibody diluent (Dako, S3022). Slides were stained overnight at 4°C in primary antibody, then stained with Alexa Fluor 488 Anti-rabbit secondary antibody (Thermo Fisher Scientific, A32731, 1:500) and Alexa fluor 594 anti-mouse secondary antibody (Thermo Fisher Scientific, A32742, 1:500) for one hour at room temperature in the dark. Slides were washed with PBS, then stained with DAPI (Invitrogen, D3571, 300 nM) at room temperature for 15 minutes in the dark. Images were acquired on the DeltaVision high resolution fluorescent microscope (GE Healthcare) equipped with SoftWork software. OPA1 intensity was analyzed in ImageJ by separating each Z-stack into the VDAC and OPA1 channel. A set threshold was applied to each channel and the images were segmented to generate regions of interest. Normalized VDAC and OPA1 intensity was calculated by dividing the area of the pixels by mean intensity for each channel. Normalized OPA1 intensity was divided by normalized VDAC intensity to generate the final OPA1 intensity/image. For immunohistochemistry (IHC), PDX slides were stained with mouse anti-human mitochondria antibody (Abcam, ab92824, 1:1000) for 20 minutes at room temperature by Agilent autostainer Link 48. Images were acquired using an Aperio digital pathology slide scanner (Leica Biosystems). Image analysis was performed with the Vectra 3 software to quantify the H-score (considering only the tumor compartment and not the stroma) for human mitochondria.

### Antibodies

The following primary antibodies were used for western blotting: anti-MFN1 (1:1000, Cell Signaling Technology, 14739S), anti-MFN2 (1:1000, Cell Signaling Technology, 9482S), anti-OPA1 (1:1000, Cell Signaling Technology, 80471S), anti-DRP1 (1:1000, Cell Signaling Technology, 8570S), anti-DRP1(S616) (1:1000, Cell Signaling Technology, 3455S), anti-FIS1(1:1000, Abcam, ab156865), anti-VDAC (1:1000, Cell Signaling Technology, 4661S), anti-β-tubulin (1:1000, Cell Signaling Technology, 2146S) and horseradish peroxidase-conjugated anti-rabbit IgG H&L (1:10,000, Abcam, ab205718).

### Seahorse

OCR and ECAR were measured with the Seahorse XFe96 extracellular flux analyzer (Agilent). Cells were seeded on XFe96 cell culture microplates and incubated at 37°C at a density of ∼1000 to 5000 cells per well 4 days prior to analysis. For the *ex vivo* seahorse assay, XFe96 cell culture microplates were coated by Cell-Tak cell adhesive solution (22.4 μg/ml, Corning # 354240) for 20minutes at room temperature and washed twice using sterile water. ∼1×1.0e^5^ purified tumor cells were seeded per each Cell-Tak coated well and then centrifugated at 200 x *g* for 1 minutes and proceed further analyses. On the day of analysis, oligomycin (1.5μM), FCCP (1μM), and rotenone A/antimycin (0.5μM) were loaded into the injection ports in the XFe96 sensor cartridge and loaded into XFe96 analyzer for calibrating. During calibration, cells were equilibrated for one hour in XF base assay medium supplemented with 10 mM glucose, 1 mM sodium pyruvate, and 2 mM L-glutamine in a CO_2_-free incubator at 37°C. After calibration, the microplate was loaded into the Seahorse bioanalyzer for analysis. Following the assay, plates were removed, then cells were washed with PBS and fixed with 3.7% formaldehyde solution by incubating for 15minutes. Cells were then permeabilized with 0.2% Triton X-100 and treated with 300nM of DAPI fluorescent dye solution in PBS (Invitrogen, D3571). DAPI-stained cells were counted on a Cytation5 (BioTek) Florescent plate reader. All OCR and ECAR data were normalized to total cell counts from DAPI staining.

### Cellular viability and growth measurement

Cell viability was measured using the CellTiter-Glo® 2.0 Cell Viability Assay kit (Promega, PR-G7572) according to the manufacturer’s instructions. Briefly, cells were seeded into 96-well culture plates at a density of ∼1000 cells/well with 100μl of growth media and incubated at 37°C. 24 hours after seeding, cells were treated with drugs at 10 different doses or IC_50_ in growth medium. 72 hours after treatment, 100ul of CellTiterGlo reagent was added to each well and incubated for 10 minutes at room temperature protected from light. Then, luminescence was measured using a FLUOstar Omega plate reader (BMG Labtech). Cell growth was assessed by measuring cell culture well confluence using an IncuCyte ZOOM (ESSEN BIOSCIENCE). Cells were seeded at 1 × 10^3^ per well in a 96-well plate, which was loaded in the IncuCyte ZOOM system. Live-cell images were obtained every 4 hours using a 10x objective lens (four images per well) within the instrument. Cell confluence was analyzed using IncuCyte image analysis software (Essen Bioscience).

### ROS Assay

The ROS-Glo™ H_2_O_2_ Assay (Promega G8820) was performed in accordance with the manufacturer’s instructions. Briefly, cells were seeded into 96-well culture plates at a density of ∼1000 cells/well with 100μl of growth media and incubated at 37°C. 48 hours after seeding, cells were treated with vehicle or drugs at IC_50_ in growth medium. 48 hours after treatment, cells were treated with H_2_O_2_ substrate solution in growth medium and incubated for 6 hours at 37°C. ROS-Glo™ detection solution was then added to each well, incubated for 20 minutes. After incubation, luminescence was measured using a FLUOstar Omega plate reader (BMG Labtech).

### *In vitro* stable isotope tracer analysis

[^13^C5]-Glutamine and [^13^C6]-glucose were from Cambridge Isotope Laboratories, Inc. (Tewksbury, MA, USA). Optima LC/MS-grade water, methanol (MeOH), ammonium hydroxide, and acetic acid (AA) were from Thermo Fisher Scientific. All other reference standards and chemicals were from Millipore Sigma. The 4-micron Dionex IonPac AG-11-HC guard column (2 × 50 mm) and 4-micron IonPac AS11-HC analytical IC column (2.0 × 250 mm) were from Thermo Fisher Scientific. The Vanquish Ultra-High Performance Liquid Chromatography system, Dionex ICS-5000+ capillary IC system, and Orbitrap Fusion Mass Spectrometer were from Thermo Fisher Scientific. To determine the incorporation of ^13^C_6_-glucose carbon and ^13^C_5_-glutamine carbon into intracellular TCA cycle and glycolysis pathway intermediates, extracts were prepared and analyzed by high-resolution, accurate-mass (HRAM)-MS. TNBC cells were seeded in 6cm petri dishes at a density of ∼2×1.0e^5^ cells in growth medium containing doxorubicin, paclitaxel, or carboplatin for 48 hours. Cells were then incubated in 2 mM of [^13^C_5_]-glutamine or 11.1 mM [^13^C_6_]-glucose RPMI media with dialyzed FBS for 48 hours before harvesting. Metabolites were extracted in 1 ml ice-cold 80/20 (v/v) methanol/water containing 0.1% ammonium hydroxide. Samples were then vortexed for 2 minutes, centrifuged at 17,000 x *g* for 5 minutes at 4°C, and supernatants were transferred to clean tubes and evaporated to dryness under nitrogen. Samples were reconstituted in deionized water, then 10μl was injected into a Thermo Scientific Dionex ICS-5000+ capillary ion chromatography (IC) system containing a Thermo IonPac AS11 250×2 mm 4 μm column. IC flow rate was 360 μl/min (at 30ºC), and the gradient conditions were as follows: started with an initial 1 mM KOH, increased to 35 mM at 25 min, then to 99 mM at 39 min, and maintained at 99 mM for 10 minutes. The total run time was 50 min. To assist desolvation and increase sensitivity, methanol was delivered by an external pump and combined with the eluent via a low dead volume mixing tee. Data were acquired using a Thermo Orbitrap Fusion Tribrid Mass Spectrometer under negative electrospray ionization (ESI) mode. The heated electrospray ionization (H-ESI) source and global MS parameters for the acquisition were as follows: scan range = 80-800 m/z; multiplex ions = false; isolation mode = quadrupole; detector type = Orbitrap; Orbitrap resolution = 240,000 (at m/z 200); RF lens (%) = 50; AGC Target = 200,000; injection ions for all available parallelizable time = true; maximum injection time (ms) = 100; microscans = 1; data type = profile; polarity = negative; source fragmentation = disabled; use EASY-IC = true; include start and end times = false; scan cycle time = 3 seconds. The raw data files were imported to Thermo Trace Finder software for final analysis. For ^13^C isotopolog analysis, ElemCor software was used for natural abundance correction and enrichment calculation^57^. The relative abundance of each isotopolog was calculated as the isotopolog area divided by (normalized by) the sum of all isotopolog peak areas.

### Statistical analysis

The data were shown as mean ± standard error (SEM), and all experiments were repeated at least three times. All statistical tests were two-sided and performed using GraphPad Prism software. All data meet normal distribution and have uniform standard deviations. Paired-sample and independent-samples were used for comparisons between two groups. For three or more groups, one-way ANOVA was used to detect the difference among groups. *P*-value less than 0.05 is considered as significant, while value less than 0.01 is considered as highly significant.

## FIGURE LEGENDS

**Supplementary Figure S1.**
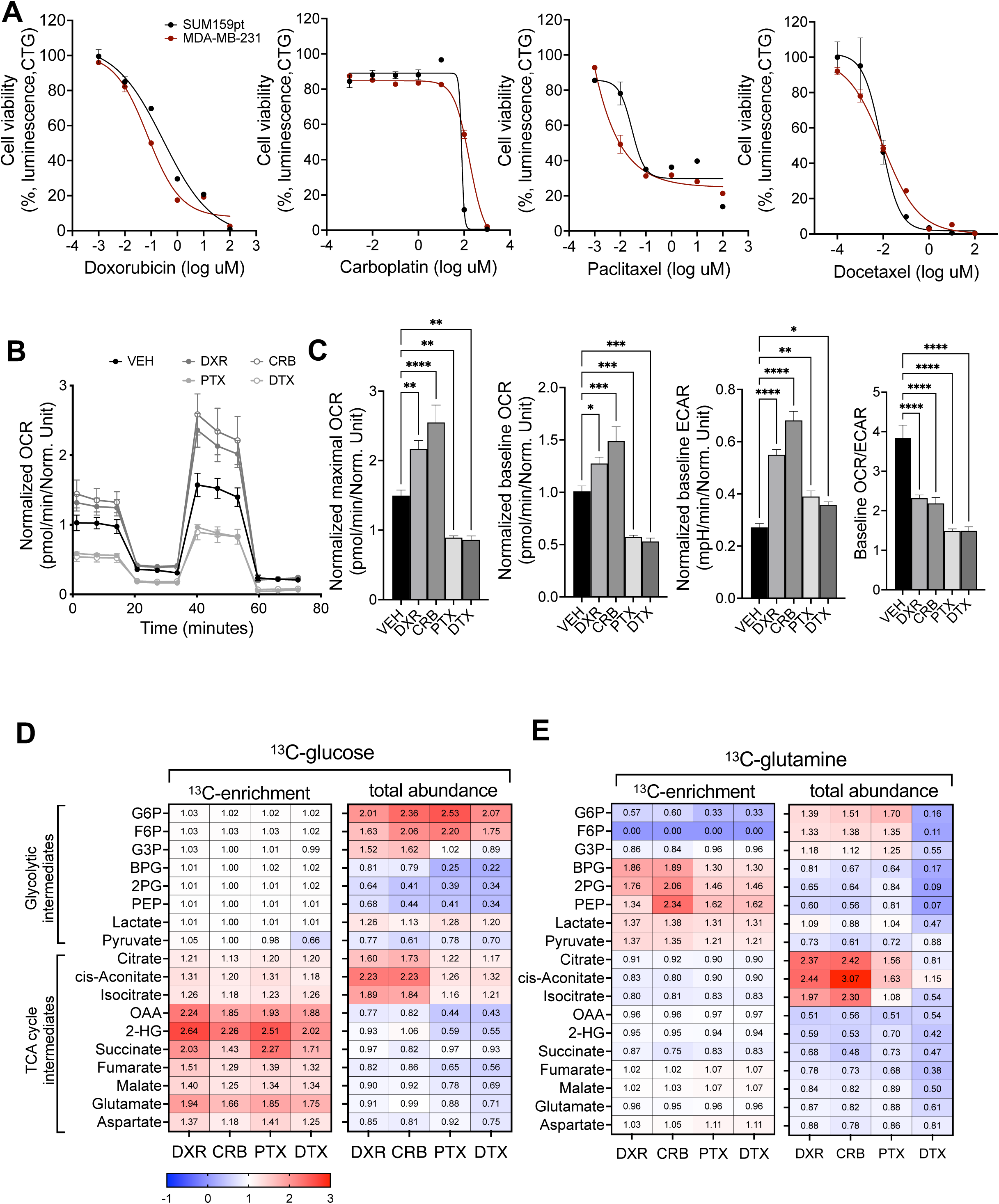
Metabolic analysis of chemotherapy treated TNBC cells. (A) Dose-response curves of chemotherapy drugs in TNBC cell lines using Cell Titer Glo Luminescence. Data are represented as the mean ± SEM (n=3). (B) Representative Seahorse Mito Stress assay of SUM159pt cells treated with vehicle or chemotherapies with (C) quantified maximal OCR, basal OCR, basal ECAR, and OCR/ECAR ratio. Data were normalized to total number of cells stained with DAPI immediately after Seahorse assay, *\*\*\*\*P* < 0.0001, ***<0.001 **<0.01 *<0.05 by one-way ANOVA. Data are representative of at least 3 independent experiments. (D-E) ^13^C-glucose derived (D) or ^13^C-glutamine (E) derived-enrichment or total abundance of glycolytic and TCA cycle intermediates in SUM159pt cells treated vehicle, DXR, CRB, PTX, or DTX. Data were normalized to vehicle treated cells. n=3, G6P (Beta-D-Glucose 6-phosphate), F6P (Fructose 6-phosphate), G3P (Glycerol 3-phosphate), BPG (2,3-Diphosphoglyceric acid), 2PG (2-Phospho-D-glyceric acid), PEP (Phosphoenolpyruvic acid), OAA (Oxoglutaric acid), 2-HG(2-Hydroxyglutarate).

**Supplementary Figure S2.**
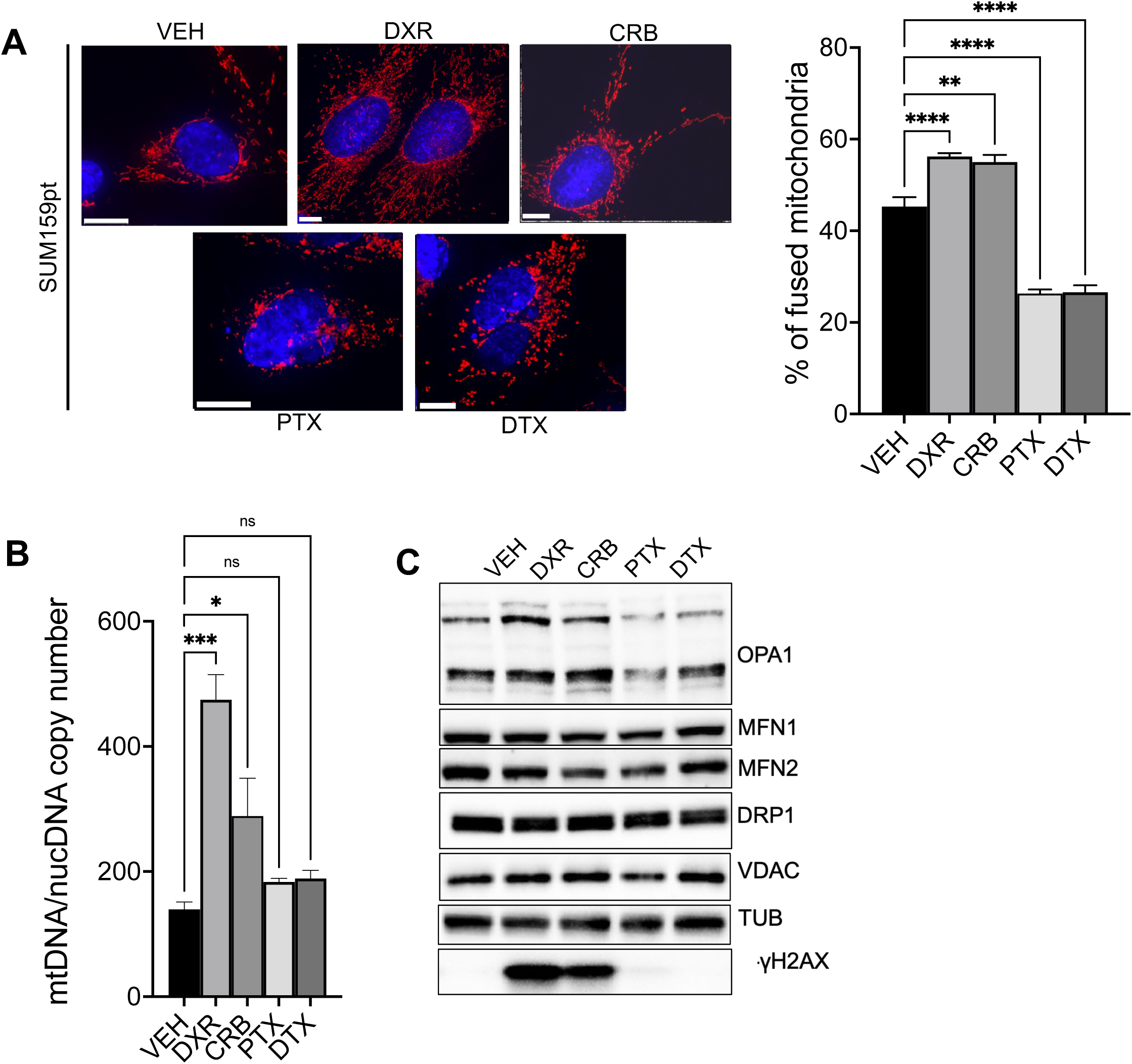
Mitochondria structure changes in SUM159pt cells. (A) Representative fluorescence images of SUM150pt cells treated with vehicle or IC_50_ of chemotherapies then stained with DAPI and MitoTracker red CMXRos. Mitochondrial morphology subtypes were quantified in at least 30 cells per experiment were analyzed, Scale bar is 10 μm. Red fluorescence, mitochondria; blue fluorescence, DAPI-labeled nucleus. (B) quantified mtDNA copy number: nuclear DNA copy number ratio, Data are represented as mean ± SEM (n=3), *\*\*P* < 0.01 by one-way ANOVA. (C) Western blot analysis of mitochondrial morphology regulators. Data are representative of at least 3 independent experiments.

**Supplementary Figure S3.**
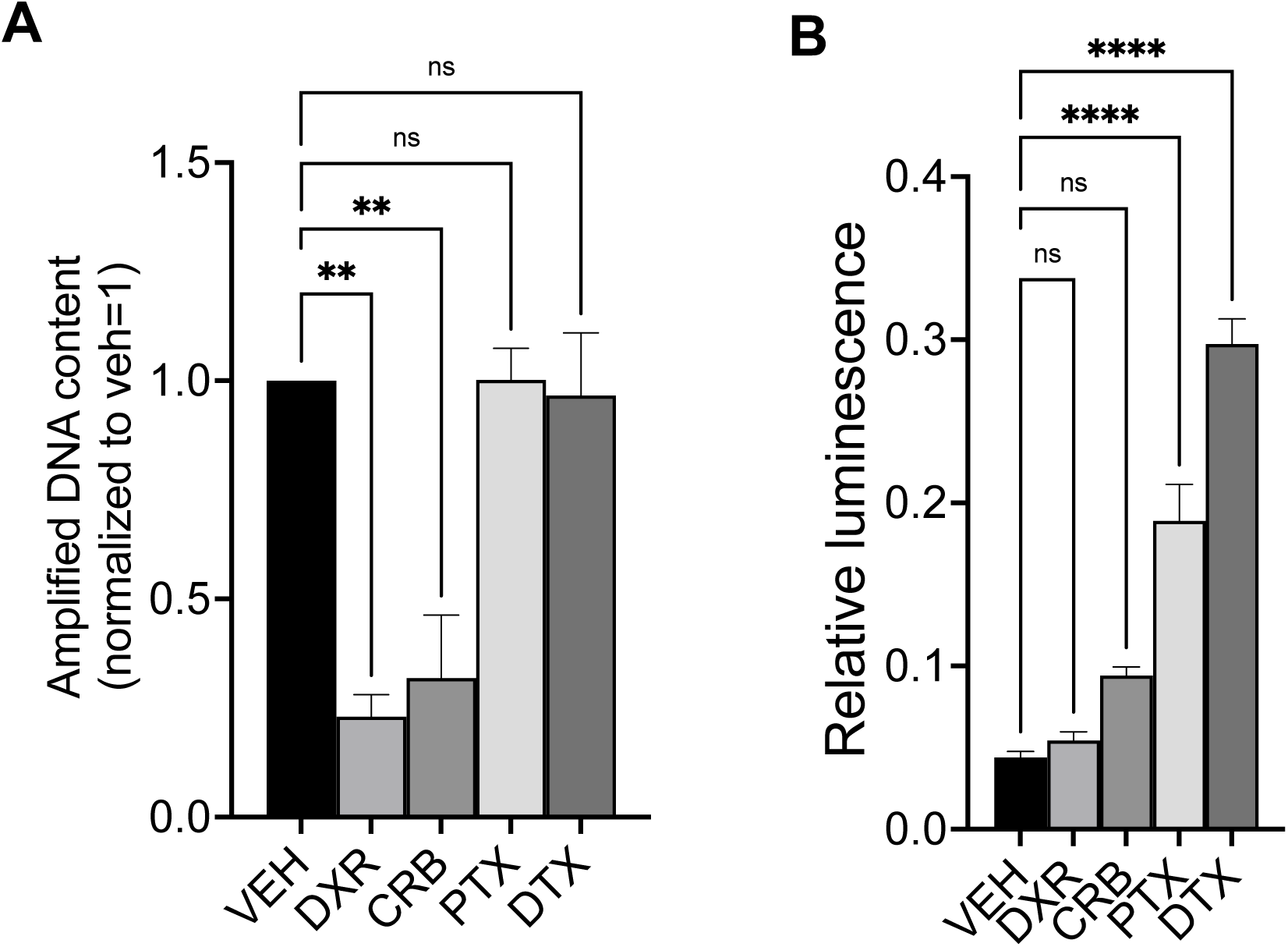
ROS and DNA damage analysis of chemotherapy-treated MDA-MB-231 cells. (A) mtDNA damage was measured in MDA-MB-231 treated different chemotherapy drugs compared to vehicle using long amplification-PCR followed by qPCR to quantify long (undamaged) mtDNA amplicons. Data are represented as mean ± SEM (n=3), *\*\*\*\*P* < 0.0001 by one-way ANOVA. (B) Results of the luminescence-based ROS assay (ROS-Glo™) of MDA-MB-231 cells. Raw luminescence values were normalized to viable cell contents that was assessed by Cell-Titer-Glo™ luminescence assay. Data are represented as mean ± SEM (n=3), *\*\*P* < 0.01 by one-way ANOVA.

**Supplementary Figure S4.**
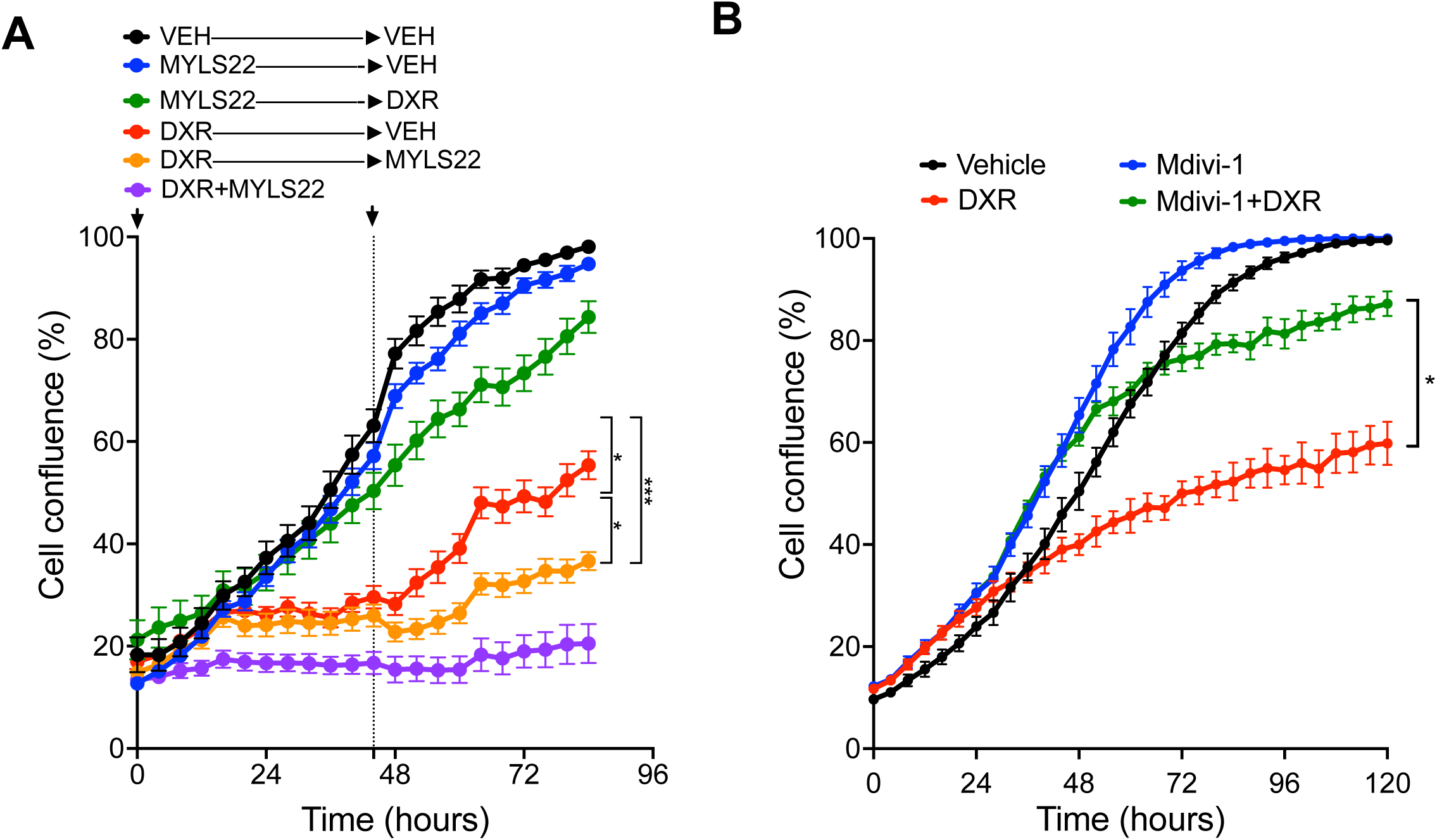
Combination treatments of MDA-MB-231 cells. (A) Representative IncuCyte-based time lapse confluence of MDA-MB-231 treated with vehicle, MYLS22 only, DXR only, MYLS22 followed by DXR, DXR followed by MYLS22, or DXR given simultaneously with MYLS22. Each data point is mean ± SD of 8 wells. (B) Representative IncuCyte time lapse confluence of MDA-MB-231 treated with vehicle, Mdivi-1 only, DXR only, or Mdivi-1 followed by DXR given after 48 hours.

**Supplementary Figure S5.**
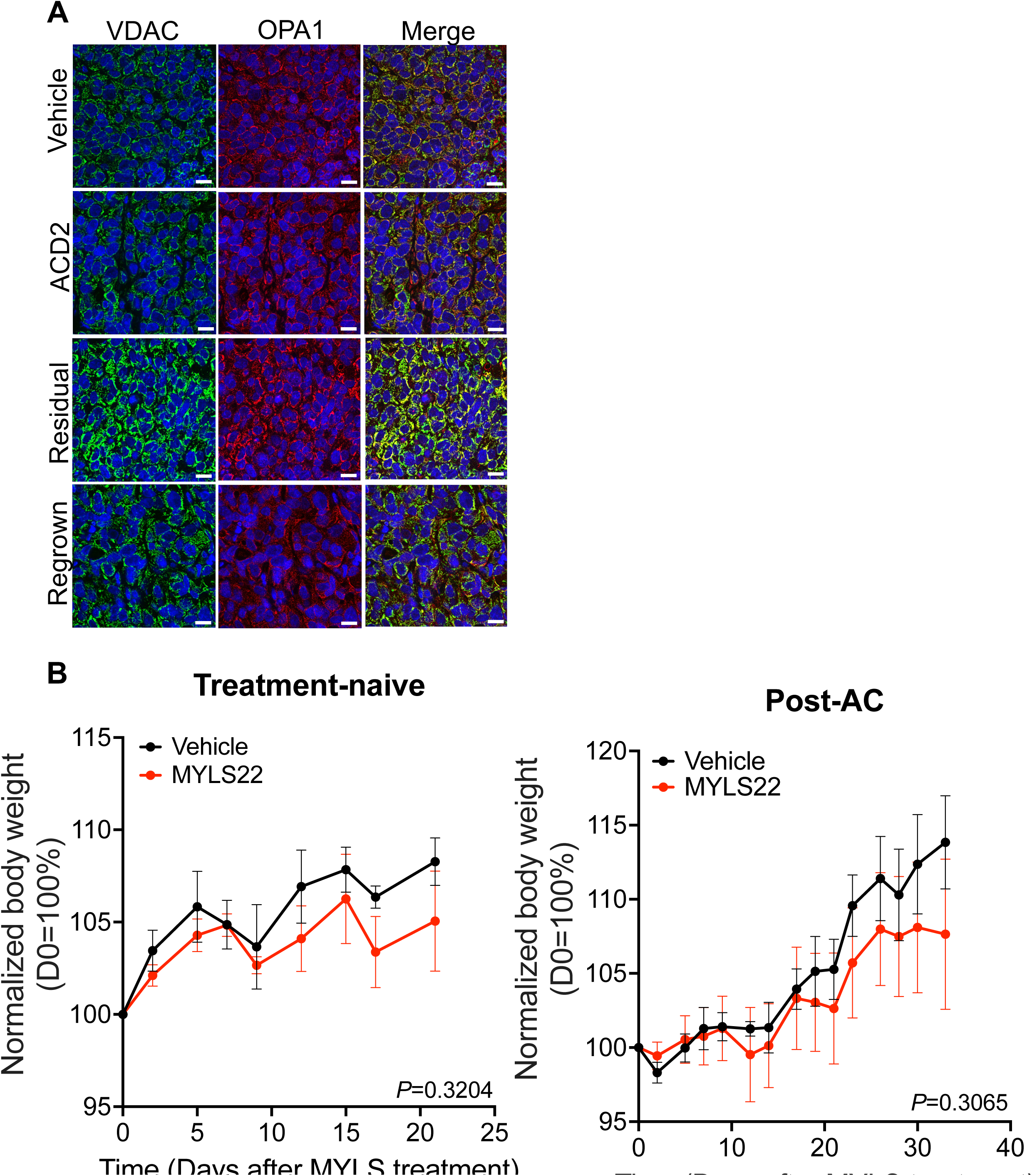
Co-immunofluorescence analysis of OPA1 and VDAC1 in PIM001-P tumor sections. (A) Representative immunofluorescence images of the tumors stained with a VDAC (green, mitochondria) antibody, OPA1 antibody (red) and nuclear dye DAPI (blue). Scale bar is 20 μm. (B) normalized body weight curve of NOD/SCID mice bearing PIM001-P tumors that were treated with an inhibitor of OPA1, MYLS22, or vehicle in the treatment-naïve (left) or in the residual setting after AC treatment (right). Error bars represent SEM (n=4–6 per group, two-sample).

## Acknowledgements

We are grateful to the breast cancer patients who donated their biopsies for cell lines and PDX models. Ms. Janice Cowden and Mr. Joshua Newby provided research advocacy support for this work. Dr. Helen Piwnica-Worms provided the PIM001-P PDX model. Fluorescence microscopy analysis was conducted at the Integrated Microscopy Core at Baylor College of Medicine and the Center for Advanced Microscopy and Image Informatics (CAMII) with funding from NIH (DK56338, CA125123, ES030285), and CPRIT (RP150578, RP170719), the Dan L. Duncan Comprehensive Cancer Center, and the John S. Dunn Gulf Coast Consortium for Chemical Genomics. Transmission electron microscopy was conducted at the Texas Children’s Hospital pathology facility. Seahorse was conducted at the Mouse Metabolism and Phenotyping Core supported by NIH UM1HG006348 and NIH R01DK114356, NIH R01HL130249. Metabolomics studies were conducted at The Metabolomics Core Facility is supported by NIH grants S10OD012304-01 and P30CA016672. STR cell line validation was conducted by the Cytogenetics and Cell Authentication Core at M.D. Anderson Cancer Center. Immunohistochemistry was conducted at the Pathology Core and Lab was created to support the research and clinical activities of the Breast Center at Baylor College of Medicine, which is supported by the Breast Center and a variety of research grants awarded to its faculty, including one of only nine Specialized Programs of Research Excellence (SPORE) in Breast Cancer granted by the National Institute of Health. Vectra microscopy and analysis was conducted with the Pathology and Histology (HTAP) at Baylor College of Medicine with funding from P30 Cancer Center Support Grant (NCI-CA125123).

## AUTHOR CONTRIBUTIONS

M.L.B. and G.V.E. were responsible for overall study design, experimentation, data interpretation, and writing of the manuscript.

J.L. conducted animal experiments and quantitative PCR under the supervision of G.V.E.

K.E.P. conducted IHC, co-IF, Vectra, and assisted with Seahorse assays under the supervision of G.V.E.

M.J.B. conducted mitochondrial assays under the supervision of G.V.E.

E.B.G. assisted with confocal microscopy image analysis under the supervision of G.V.E.

L.T. and S.A.M. conducted metabolomics sample processing and analysis under the supervision of P.L.L.

T.W. conducted statistical analyses.

M.M. conducted TEM of TNBC cell lines.

B.L. assisted with analysis of chemotherapy experiments.

J.P.B. conducted TEM of PDX tumor tissues and assisted with analysis.

W.P. assisted with analysis of TEM and mitochondria function experiments.

P.L.L. oversaw metabolomics experiments and analyses.

All authors have critically read, edited, and approved the final version of this manuscript.

